# Bone Marrow Spatial Transcriptomics Reveals a Myeloma Cell Architecture with Dysfunctional T-Cell Distribution, Neutrophil Traps, and Inflammatory Signaling

**DOI:** 10.1101/2024.07.03.601833

**Authors:** Laura Sudupe, Emma Muiños-Lopez, Ana Rosa Lopez-Perez, Amaia Vilas-Zornoza, Sarai Sarvide, Purificacion Ripalda-Cemborain, Paula Aguirre-Ruiz, Patxi San Martin-Uriz, Marta Larrayoz, Laura Alvarez-Gigli, Marta Abengozar-Muela, Itziar Cenzano, Miguel Cócera, Javier Ruiz, Ignacio Sancho González, Azari Bantan, Aleksandra Kurowska, Jin Ye, Phillip T. Newton, Bruno Paiva, Juan R. Rodriguez-Madoz, Vincenzo Lagani, Jesper Tegner, Borja Saez, Jose Angel Martinez-Climent, Isabel A. Calvo, David Gomez-Cabrero, Felipe Prosper

## Abstract

The bone marrow (BM) is a complex tissue where spatial relationships influence cell behavior, signaling, and function. Consequently, understanding the whole dynamics of cellular interactions requires complementary spatial techniques that preserve and map the architecture of cell populations *in situ*. We successfully conducted spatial transcriptional profiling using Visium Spatial Gene Expression in formalin-fixed paraffin-embedded (FFPE) BM samples obtained from healthy and Multiple Myeloma (MM) mouse models and patients, addressing the technical challenges of applying spatial technology to long bone samples. A custom data-analysis framework that combines spatial with single-cell transcriptomic profiles identified both the BM cellular composition and the existing cell relations. This allowed us to visualize the spatial distribution of transcriptionally heterogeneous MM plasma cells (MM-PC). We spatially delineated transcriptional programs associated with MM, including NETosis and IL-17-driven inflammatory signaling, which were inversely correlated to malignant PC-enriched regions. Furthermore, a gradient of MM-PC density spatially correlated with a shift from effector-to-exhausted T cell phenotypes. The translational relevance of our findings was confirmed using FFPE BM biopsies from MM patients with varying levels of malignant PC infiltration. In summary, we provide the first spatial transcriptomics analysis applied to a mouse and human mineralized bone tissue and illustrate the BM cellular architecture of MM, revealing deregulated mechanisms underlying MM intercellular communication.

## INTRODUCTION

The bone marrow (BM) microenvironment is a specialized niche with stromal cells, blood vessels, immune cells, and extracellular matrix components, crucial for regulating hematopoietic stem cells (HSC). This environment supports cell growth, differentiation, and migration, impacting both normal physiology and disease^1^. The role of the BM microenvironment in the development and progression of Multiple Myeloma (MM), a B-cell malignancy characterized by the clonal expansion of abnormal plasma cells (PC) that infiltrate the BM^2^, has clearly been demonstrated^3–7^. Various mechanisms have been implicated in the expansion and growth of malignant PC. For example, the release of matrix metalloproteinases (MMPs) facilitates invasion into nearby tissues and metastasis, or the secretion of interleukins promotes myeloma cell growth or inhibits immune function^6,8^. Moreover, the induction of neutrophil extracellular traps (NETs) formation by neutrophils, or the impairment of T cell function, allows tumor expansion^7,9^. All these pieces of evidence suggest that understanding the interplay between malignant cells and their BM microenvironment may contribute to the identification of novel targeted therapies^10,11^. How these interactions are geographically distributed within the BM, the different relations between multiple myeloma PC (MM-PC) and other cellular components, and the impact that these interactions have on transcriptional programs is still unclear.

Advances in spatial transcriptomics have provided the means to establish unbiased gene expression analysis with spatial context for different tissues, making these technologies complementary to single-cell methods that lack spatial resolution^12–14^. While tissues such as the heart or brain have been recently characterized through spatial transcriptomics, mineralized tissues such as BM possess a particular challenge^15–17^. Recent publications overcoming the spatial limitations of mineralized bone tissue^18,19^, combined with the integration of various predictive modeling packages for long bones^16^, have led to significant advancements in studying BM regulation. However, the complexity of MM pathology and the use of healthy and malignant human samples continue to present significant challenges^20^.

In the current study, we have used Visium Spatial Gene Expression (10x Genomics) technology on formalin-fixed paraffin-embedded (FFPE) bone femur sections of our recently described healthy and MI_cγ1_ MM mouse models^21^, as well as in human FFPE-BM samples from healthy and diseased individuals. This approach allowed us to explore the spatial organization of the BM in homeostasis and MM, while elucidating the specific transcriptional regulatory programs of malignant PC and their surrounding microenvironment. Using a custom data-driven approach, we defined specific areas of MM-PC infiltration, allowing examination of the surrounding immune cells and their interactions based on the concentration of tumor cells. Our results also highlight MM-PC transcriptional heterogeneity between different malignant-cell-enriched areas. We spatially defined transcriptional programs potentially involved in the pathogenesis of MM within the BM microenvironment that were associated with MM-PC density gradient. Among them, we observed the spatial location of T cells with effector and exhausted phenotypes, the gradual decrease in inflammatory pathways, and the formation of NETs by neutrophils in areas of MM infiltration. Importantly, the identification of different areas depending on MM-PC infiltration and the interaction between malignant cells and their microenvironment were confirmed in BM biopsies from MM patients with varying degrees of tumor PC infiltration. Our study introduces a novel approach to understanding MM by identifying the spatial interactions between MM-PC and their BM microenvironment, which may prove key for identifying resistance mechanisms.

## RESULTS

### Spatial resolution of healthy bone marrow resident cell types

To elucidate the complex interactions between multiple MM-PC and their surrounding microenvironment, we analyzed the spatial molecular landscape of cells within both healthy (Fig. 1 and 5) and malignant (Fig. 2-5) BM tissue. Building on previous reports^16^, we first performed spatial transcriptomics of mouse femurs from a preclinical healthy mouse model^21^ (YFPcγ1) using Visium Gene Expression (10X Genomics) (Fig. 1a and Table S1). After preprocessing and quality control analysis, we detected a total of 1242 spots with an average of 3998 features and 14118 counts per spot in the trabecular region of the bone (Fig. 1b). The cortical region presented a minimal number of counts (data not shown), and the cells in the fully mineralized bone did not interact directly with the disease-associated cells, leading to its exclusion from the analysis.

**Fig 1.**
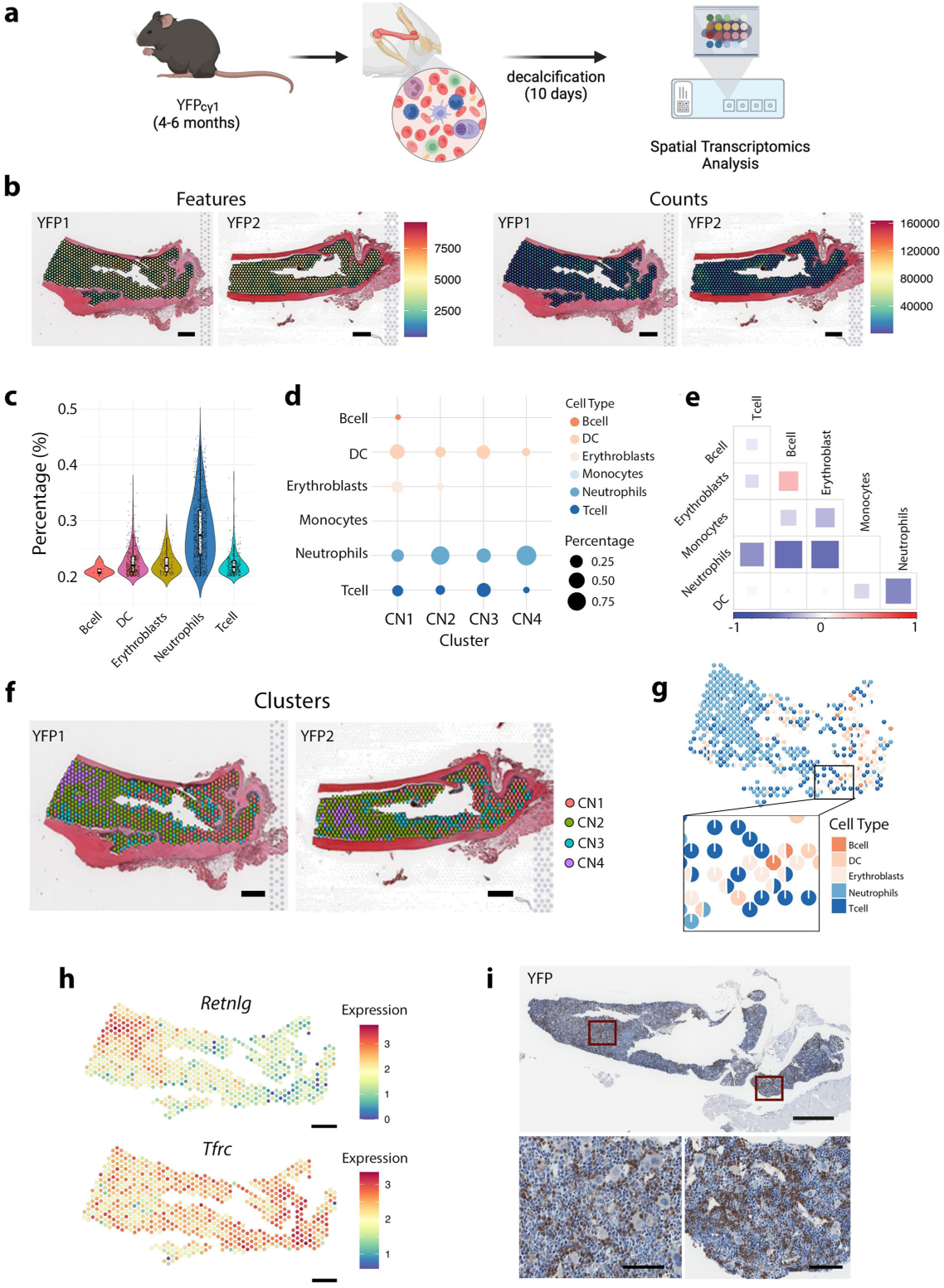
Spatial resolution of healthy bone marrow resident cell types. **a** Schematic representation of healthy control mouse (YFPcγ1), experimental workflow for bone tissue isolation and spatial transcriptomics analysis. **b** Quality control of the mice femur tissues, Features, and Counts in two samples from a control mouse model (YFP1 and YFP2)^16^. **c** Cell type proportions per spot derived from a deconvolution analysis using scRNA-seq data as a reference^19^. **d** Average of the most abundant cell types in the four identified clusters (CN1-CN4). **e** Correlation analysis of cell types based on spot distribution. **f** Spatial distribution of the four identified clusters. **g** Pie charts illustrating each cell type’s proportion to each spot’s transcriptomic signature. **h** *Retnlg* and *Tfrc* mouse gene expression, canonical markers of neutrophils and erythroblasts, respectively. **i.** Immunohistochemistry (IHC) of Ly-6G/Ly-6C (*Gr-1*), a neutrophil marker, in control mice femur sample (YFP). Scales of 500 μm (b, f, and h), scales of 1000 μm (upper panel i) and 50 μm (bottom panels i).

**Fig 2.**
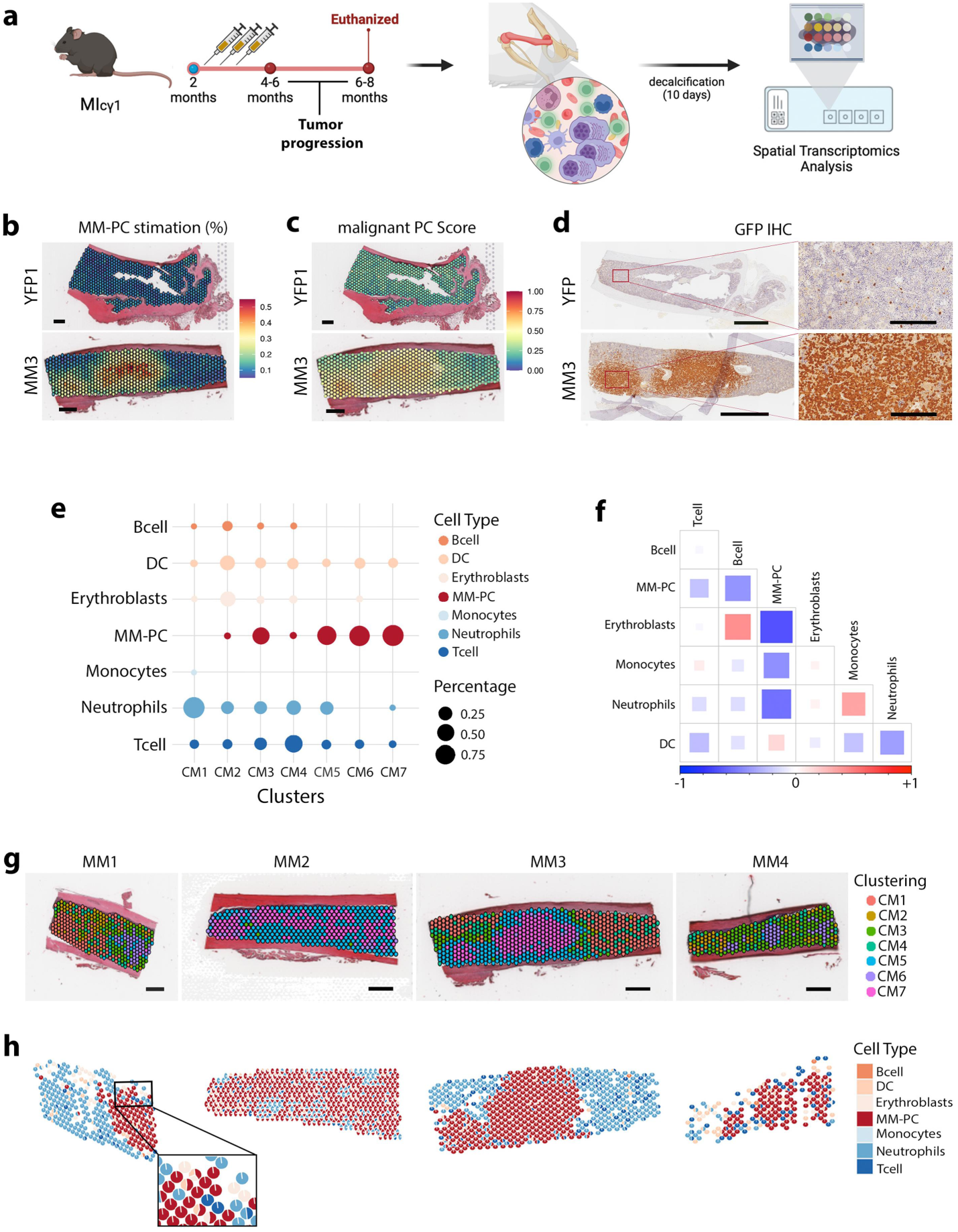
Unraveling the transcriptional spatial profiling of multiple myeloma bone marrow. **a** Schematic representation of MM mouse model (MIcγ1), experimental workflow for isolation bone tissue and conducting spatial transcriptomics. **b** Estimating malignant plasma cell (MM-PC) percentage per spot by deconvolution analysis using the scBMReference data set. **c** enrichment score for malignant PC. **d** Green Fluorescent Protein (GFP) IHC of healthy YFPcγ1 (YFP1) and pathological MIcγ1 mice (MM3) femur tissues. **e** Average of the most abundant cell types in each of the seven identified malignant clusters (CM1-CM7). **f** Spot-based cell type correlation analysis between the different cell types. **g** Spatial distribution of the clusters identified in malignant samples. **h** Pie charts representing each cell type’s contribution to each spot’s transcriptomic signature. Scales of 500 μm (b, c, and g) and scales of 1000 μm (left panels d) and 100 μm (right panels d).

Since the BM is an unshaped tissue and each spatial transcriptomic spot captured a group of cells (between 3 and 10 cells)^22^, we estimated the cell composition per spot. To this end, based on the literature, we first identified the most prevalent cell types in the BM^23^. Secondly, we estimated the cell proportion per spot by deconvolution analysis using single-cell RNA-seq (scRNA-seq) data as a reference^24^ (Fig. 1c and Fig. S1a). Our spatial transcriptomics revealed neutrophils as the major cell type population, consistent with previous reports^23^. Next, we conducted a clustering analysis using the estimated cell proportion. We identified four groups of spots (CN1-CN4) (Fig. 1d). “Major cell type per spot” analysis (see Methods) was performed to identify the dominant cell type influence. Clusters 1 (CN1), 2 (CN2), and 3 (CN3) were characterized by a larger proportion - when comparing between clusters - of DC, neutrophils, and T cells, respectively. Cluster 4 (CN4) exhibited the highest estimated percentage of neutrophils, while erythroblasts and B cells were reduced in this cluster. We observed a positive spatial correlation between erythroblasts and B cells, suggesting that these two cell types may establish functional interactions (Fig. 1e). In contrast, neutrophils presented negative correlations with erythroblasts, B cells, DC, and T cells, indicating a spatial distinction where neutrophils are less prevalent in regions where erythroblasts, B cells, DC, and T cells are more abundant.

The spatial distribution analysis of the different clusters revealed differences between the diaphysis (central area of the femur) and the epiphysis (extremes of the femur) (Fig. 1f). This is consistent with the heterogeneity observed in the Hematoxylin-Eosin staining (H&E) with the Green Fluorescent Protein (GFP) immunostaining performed to identify healthy PC (visualized with Diaminobenzidine- DAB) (Fig. S1b). Notably, the spots in the healthy tissue’s diaphysis were enriched in neutrophils. In contrast, the epiphysis of the femur showed an increased presence of erythroblast (Fig. 1g). These results were consistent with the expression of the Retnlg, as well as Tfrc, canonical markers of neutrophils and erythroblasts respectively (Fig. 1h). The spatial enrichment of neutrophils in diaphysis area was further confirmed by immunohistochemistry (IHC) of Ly-6G/Ly-6C (Gr-1) (Fig. 1i). Taken together, we successfully describe the spatial configuration of the most prevalent cell type populations in healthy BM space in bone tissue.

### Unraveling the transcriptional spatial profiling of multiple myeloma bone marrow

Following the same approach, we next aimed to characterize the structure of the BM under pathological conditions, and thus, we focused on MM as a case study of a tumor characterized by BM involvement. To that end, we used our recently described MM model (MI_cγ1_), which recapitulates the principal clinical, genetic, and immunological characteristics of MYC and IKK2^NF-κB^ -driven MM patients^21^ (Fig. 2a). We spatially profiled mouse femurs from animals with different percentages of MM-PC infiltration (Table S1 and Fig. S2a) and detected a total of 1836 spatially defined spots with an average of 4914 features and 25646 counts per spot (Fig. S2b).

As myeloma samples are characterized by significant MM-PC infiltration, deconvolution of spatial spots was conducted using scBMReference (murine scRNA-seq dataset^24^, used to annotate healthy BM, combined with our MM-PC scRNA-seq^25^). As expected, tumor PC were not detected in healthy BM samples (Fig. 2b, upper panel). In contrast, we observed a large proportion of the estimated percentage of MM-PC in several locations within each sample from MM mice (Fig. 2b, bottom panel, and Fig. S2c). Next, we compared the estimated MM-PC cell proportion per spot against a malignant PC gene-set score (Table S2). This score is a unitless measure that calculates the activity of a gene-set (also referred to as a signature) per spot. We computed a malignant PC signature by comparing transcriptionally malignant PC with the cell types described at the single-cell level (Fig. 2c and Fig. S2d). As anticipated, we detected a strong correlation between the MM-PC proportion and the enrichment score for malignant PC (Fig. S2e). Based on this observation, we used cell proportions and/or scores depending on the context in the subsequent analyses. Interestingly, we observed heterogeneity in the distribution of the estimated percentage of MM-PC among the samples (Fig. S2f). This result was confirmed by the expression of GFP using IHC, as MM-PC of this model constitutively expresses GFP (Fig. 2d and Fig. S2g).

We applied the same strategy used for healthy samples to samples with MM and identified seven clusters (CM1-CM7) (Fig. S3). In clusters with a low proportion of MM-PC, clusters 1 (CM1) and 2 (CM2), we observed similar prevalence distributions to those in healthy samples, that is, the dominance of neutrophils or erythroblasts. However, the “Major cell type per spot” analysis indicated a steady proportion of T cells and DC in all the clusters, with a gradual increase in MM-PC in different clusters, revealing the maximum malignant PC proportion from clusters 5 (CM5) to 7 (CM7) (Fig. 2e). When considering all spots in the BM area, we identified a negative correlation in cell proportion between MM-PC with erythroblasts, neutrophils, monocytes, and B cells, consistent with prior studies indicating that MM-PC displaced other hematopoietic populations in the BM of MM patients^26,27^ (Fig. 2f). On the contrary, a positive correlation was seen with DC, also consistent with previous studies^28^. No specific pattern of spot distribution was identified, consistent with the described spatial clonal architecture and heterogeneous distribution of MM-PC^29^ (Fig. 2g and Fig. S3). However, identifying the different clusters allowed us to define the specific cell types surrounding the malignant PC (Fig. 2h), establishing the spatial relation between the different BM cell types and the MM-PC. Overall, these results demonstrate the feasibility of applying spatial transcriptomics to fully mineralized murine-diseased tissue and provide a comprehensive spatial overview of malignant PC heterogeneity in MM.

### Molecular characterization of plasma cell spatial heterogeneity

Looking into the spatial expression of several myeloma PC canonical markers, including Sdc1, Tnfrsf17, Mzb1, and Xpb1, we could confirm the presence of MM-PC groups in our malignant BM samples (Fig. 3a). We noted significant spatial heterogeneity in the expression of certain markers of PC within the MM-PC enriched areas, such as Mzb1 and Xbp1 (Fig. 3a and Fig. S4a-b). To further characterize this spatial transcriptional heterogeneity of malignant PC, we performed a data-driven identification of areas with high concentrations of malignant PC using the healthy samples and the malignant PC score to establish criteria (Fig. S5a). These criteria facilitated the identification of spots resembling healthy tissue, which we labeled “Rest” (areas without malignant PC). Subsequently, we classified the “non-Rest” spots into three categories—"Remote zone," "Border zone," and "Hotspot"—based on the density of MM-PC in each spot determined by the malignant PC signature (Fig. 3b). Here, given the previously observed strong correlation between malignant PC score and MM-PC proportion, "Hotspot" indicated spatial areas with high malignant PC proportion. In contrast, the "Remote zone" signified areas with low proportions, and the "Border zone" encompassed the regions surrounding the hotspot. To further understand the heterogeneity of the MM-PC, we grouped the “Hotspots” based on their spatial positions using graph-based clustering for each sample (Fig. S5b-c and see also Methods). This resulted in the identification of twelve transcriptionally distinct MM-PC groups (Pg1-Pg12) within the “Hotspot” areas among all the diseased samples (Fig. 3c). The areas and MM-PC groups identification were conducted separately for each sample to make the identified areas comparable despite the differences in MM-PC infiltration.

**Fig 3.**
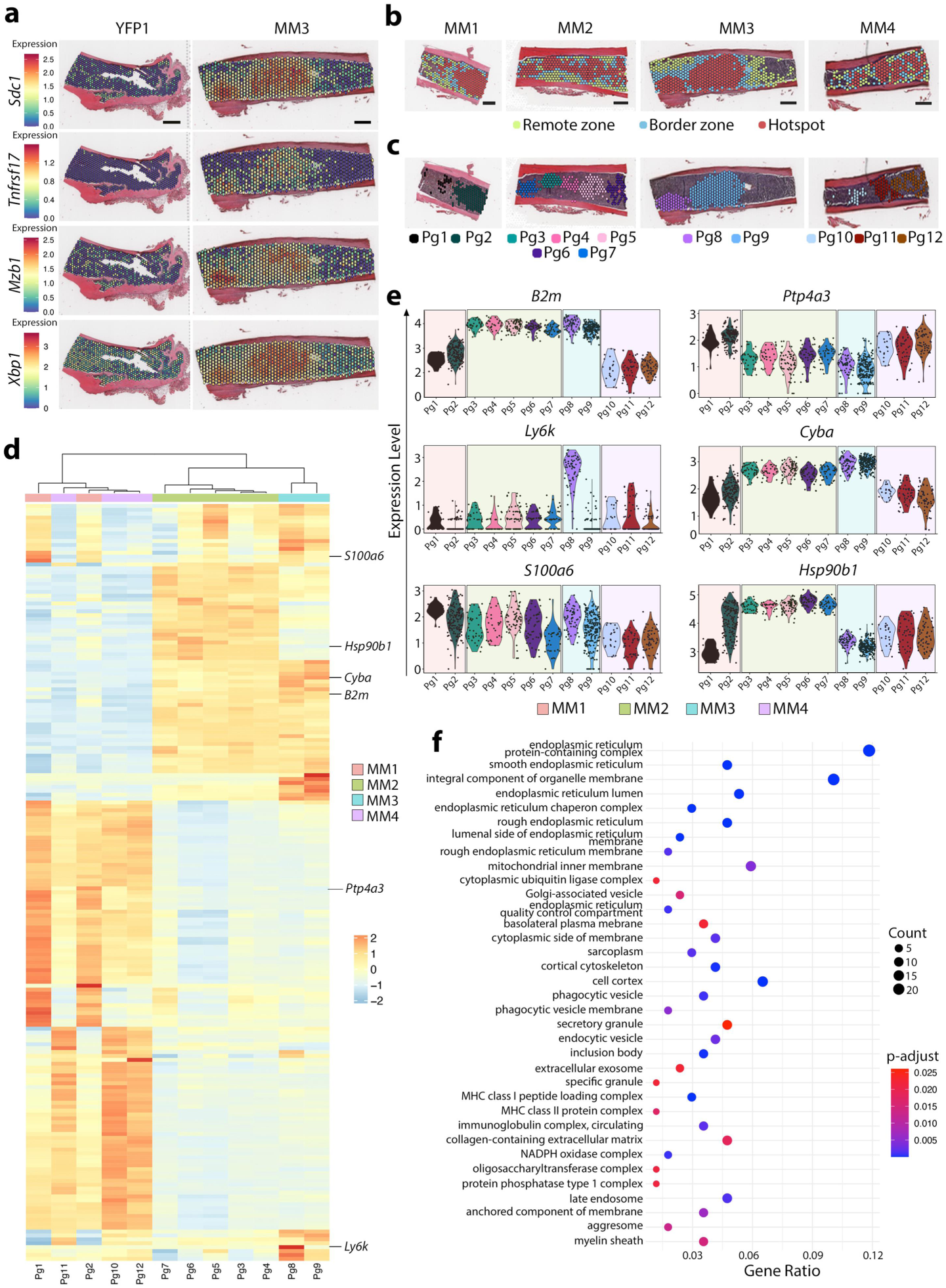
Molecular characterization of plasma cell spatial heterogeneity. **a** Spatial gene expression of pathological PC canonical markers. **b** Anatomical position of the identified areas (Remote zone, Border zone and Hotspot) based on their malignant PC signature. **c** Spatial distribution of the twelve identified MM-PC groups (Pg1-Pg12). **d** Heatmap of the top 1% most highly variable genes between the Pg groups shown in c. **e** Violin plots of selected genes from (d). **f** Selected Gene Ontology (GO) terms derived from the over-representation analysis (ORA) of the most variable genes identified in (d) across Pg groups (the entire list is in Supplementary Data 2). Scales of 500 μm (a and b).

After identifying the multiple spatial MM-PC groups, we delved into their transcriptional profiles. First, we noted a distinct separation among certain Pg, highlighting the transcriptional heterogeneity of these groups within the BM when employing pseudo-bulk profiles computed for each Pg (Fig. S5d). Secondly, we identified the top 1% of the most variable genes across the Pg1-Pg12 (n=194) (Fig. S5e and Supplementary Data 1). As a result, we observed specific expression patterns in the different MM-PC groups (Fig. 3d). Among the most variable genes, we found B2m, Ptp4a3, Ly6k, Cyba, S100a6 and Hsp90b1 (Fig. 3e and Fig. S6). These genes have been implicated in numerous physiological processes and tumorigenesis, including the development of MM^30–34^. Over- representation analysis (ORA) on the highly variable genes identified pathways involved in major histocompatibility complex (MHC), immunoglobulin complex, cytoskeleton, vesicle, extracellular matrix, and NADPH oxidase complex (Fig. 3f and Supplementary Data 2). This analysis also highlighted the endoplasmic reticulum (ER) related pathways corroborating the described role of ER and ER stress in MM’s pathogenesis and drug resistance^35,36^. In summary, these findings suggest transcriptional heterogeneity within MM and delineate distinct spatial expression patterns of known and novel genes associated with MM pathogenesis.

### Spatial identification of transcriptional programs associated with MM pathogenesis

Recognizing the BM microenvironment as a crucial contributor to the MM pathogenesis and the role of T cells in the immune response to malignant PC infiltration, we analyzed different transcriptionally defined T cells according to their spatial relation to MM-PC^3,37,38^. The score-based analysis of spatial transcriptomic signatures indicated the presence of effector T cells within the MM-PC Hotspot area (spatial region with high concentrations of malignant PC), suggesting active immune aggression potentially targeting tumor cells (Fig. 4a-c, Fig. S7a, and Supplementary Data 3). Exhausted T cells^39^ were predominant in the Border zone (areas surrounding the hotspots with less malignant PC proportion), representing a depleted immune microenvironment.

**Fig 4.**
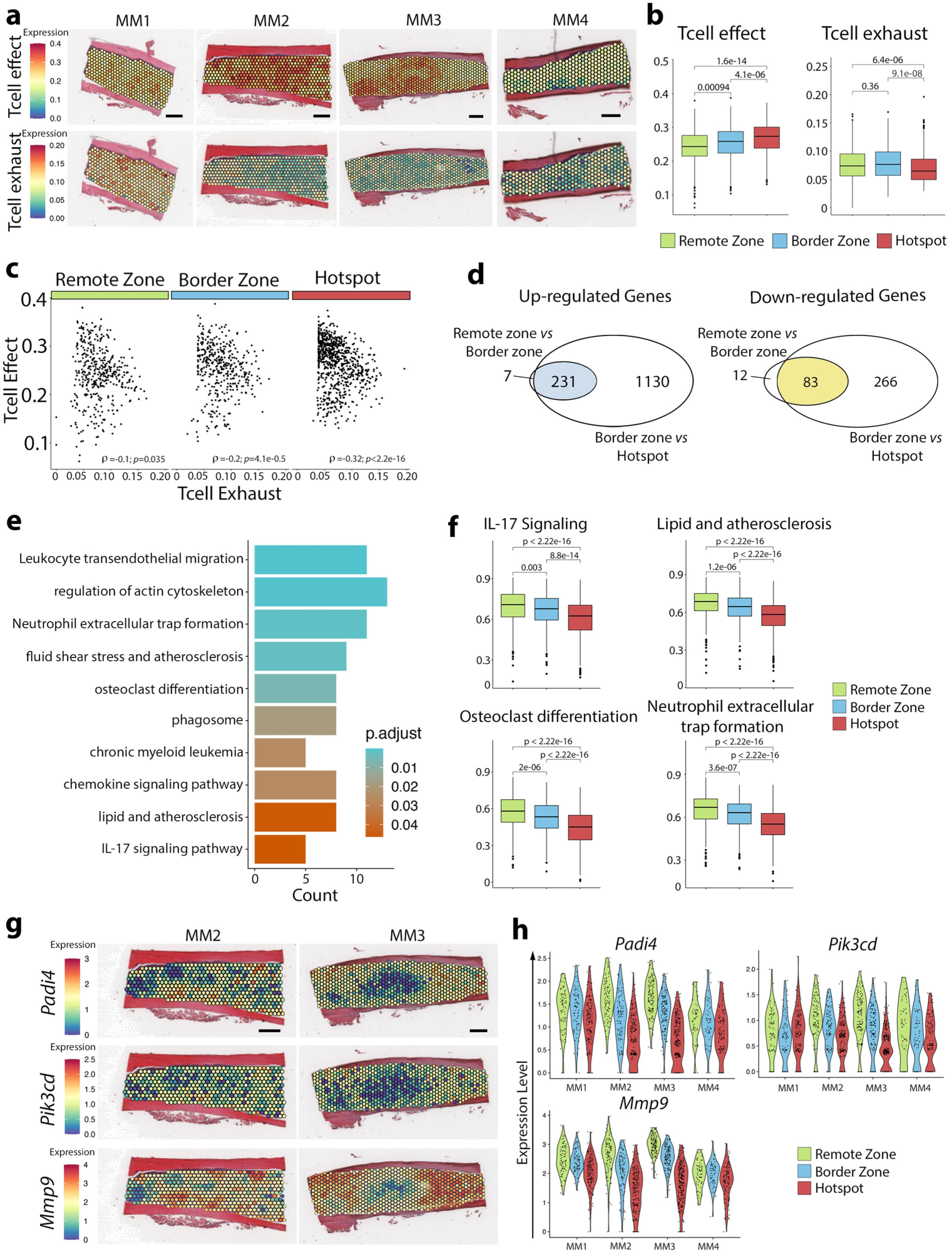
Spatial identification of transcriptional programs associated with MM pathogenesis. **a** Spatial distribution of the T cell effector (upper panels) and T cell exhaustion (bottom panels) scores in the pathological samples. **b** Box plots illustrating the T cell effector and T cell exhaustion scores across different identified areas, with their corresponding p-values. **c** Correlation between the T cell effector and T cell exhaustion profiles in the three identified areas. **d** Venn diagram displaying the differentially expressed genes (DEG) between the different areas (Remote zone, Border zone, and Hotspot). **e** Bar plot indicating upregulated KEGG pathways from the 231 commonly upregulated genes identified in (d). **f** Box plots of selected KEGG pathways from (e). **g** Spatial distribution and **h** violin plots of *Padi4*, *Pik3cd*, and *Mmp9* genes expression level. Scales of 500 μm (a and g).

Secondly, we conducted a data-driven approach to uncover additional microenvironmental signatures associated with MM. A differential expression analysis between the different areas identified 333 differentially expressed genes (DEG) between the Remote zone and Border zone (238 up- and 95 down-regulated) and 1710 DEG between the Border zone and Hotspot (1361 up- and 349 down- regulated) (Fig. 4d, Fig. S7b, and Supplementary Data 4). Focusing on the 231 commonly up- regulated genes in these comparisons, we found an enrichment in many diverse biological processes highlighting leukocyte transendothelial migration and NETs formation, among others (Fig. 4e and Supplementary Data 4). Interestingly, when investigating the associated signatures, we observed a statistically significant decreased score for pathways associated with interleukin-17 (IL-17) signaling, atherosclerosis, osteoclast differentiation, and NETosis within the Hotspots of tumor cells. These results suggested a higher activity of these pathways in the MM-PC microenvironment compared to areas with tumor cells, specifically in the Remote zone (areas with low proportions of MM-PC) (Fig. 4f). Additionally, we observed similar trends in other pathways indicative of tissue remodeling post- MM development (Fig. S7c). To gain deeper insights into the molecular mechanisms driving the MM-PC microenvironment, we analyzed the spatial distribution of highly up-regulated genes in the Remote zone, such as Padi4, Pik3cd, and Mmp9, along with Ncf4 and S100a8 (Fig 4g-h and Fig. S7d-f). Altogether, these changes point towards an impact of MM-PC infiltration on the response of the BM microenvironment against tumor cells, with a progressive decrease in inflammation, neutrophil response, and matrix remodeling in areas with a high MM-PC infiltration.

### Spatial transcriptomic analysis of human BM samples

To validate the findings obtained from mouse models, we conducted spatial transcriptomics on FFPE- BM samples from three healthy individuals and six MM patients (Fig. 5a). The clinical characteristics of these subjects are detailed in Table S3. First, we confirmed the presence of tumor regions in the samples through a comparison of H&E staining, malignant PC score, and the expression of key MM genes such as CD81, XBP1, and TNFRSF17 (Fig. 5b-d, Fig. S8a-c and Table S4). We also demonstrated that these areas overlapped with tumor regions by CD38 or CD138 IHC (Fig. 5e and Fig. S8d).

**Fig 5.**
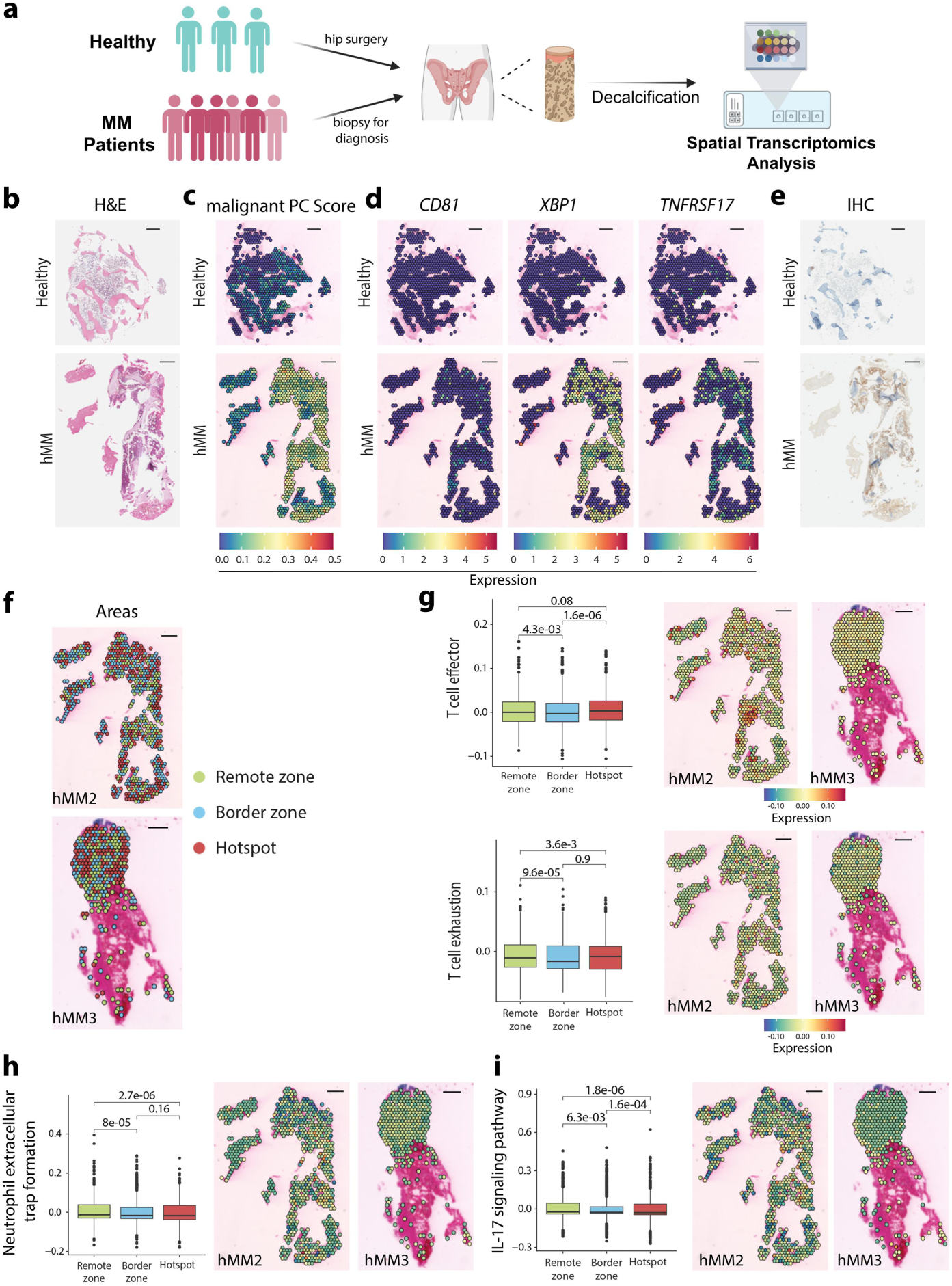
Spatial transcriptomic analysis of human BM samples. **a** Experimental workflow for isolation and spatial transcriptomics analysis of healthy and MM human bone tissue samples. **b** Hematoxylin-Eosin staining (H&E) staining. **c** spatial representation of the malignant PC enrichment score. **d** spatial gene expression of human PC canonical markers. **e** CD138 (healthy) and CD38 (human malignant bone tissue (hMM)) IHC, of healthy (upper panels) and human malignant bone tissues (bottom panels). **f** Anatomical position of the identified areas (Remote zone, Border zone, and Hotspot) in two diseases samples, hMM2 and hMM3. **g** Box plots per area (mean of all the diseased samples) and spatial distribution of the T cell effector and T cell exhaustion scores. **h,i** Box plots per area (left panel) (mean of all the diseased samples) and spatial distribution of the scores associated with Neutrophil extracellular trap formation (NETs) (**h**) and with IL-17 signaling pathways (**i**) in two human malignant tissues (hMM2 and hMM3) (middle and right panel). Scales of 500 μm (b-i).

Next, using the MM-PC signature-derived scores, we categorized the spots of human MM samples into three distinct areas: “Remote Zone,” “Border Zone,” and “Hotspot” (Fig. 5f and Fig. S9a-c) by adapting the analytical pipeline previously described for mice. The human BM’s enrichment of exhausted and effector T cells was less evident in human than in mice (Fig. 5g and Fig. S9d, e). While the effector T cells profile globally followed the same tendency between Hotspot (spatial areas with high tumor cell proportion) and surrounding areas (Border and Remote zone, areas with less proportion of MM-PC) as in mice, showing an increased score in Hotspot area, there was a slight tendency for an increase in the score of exhausted T cells also within areas with high proportion of malignant PC (Hotspot). These results suggest an increase in immune deregulation in MM patients. When we analyzed transcriptional signatures associated with NETosis and IL-17 signaling, among others, we observed a decrease in the associated scores from the Remote zone to the Hotspot area, consistent with our previous observations in mice (Fig. 5h, i and Fig. S9f-h and Fig. S10). These data suggested that these pathways were more active in the Remote zones in comparison with the Hotspot areas. Taken together these results in human samples validate our spatial transcriptomic approach and underscores its potential for clinical translation, enhancing our understanding of the MM microenvironment’s complexity and offering a robust framework for future therapeutic exploration.

## DISCUSSION

Recent studies have demonstrated the possibility of resolving the transcriptional spatial complexity of soft tissues, contributing to identifying novel mechanisms involved in tissue damage, cancer development, or inflammatory response. While this technique is readily applied to soft tissues, the requirement for bone decalcification -a process that typically degrades mRNA- has presented a significant challenge for mineralized tissues and BM samples^15,16,20^. Recent studies have demonstrated, for the first time, the potential of this technology to identify specific transcriptional pathways involved in the regulation of skeletal stem and progenitor cells in mice^16,40^ and human osteosarcomas^41^. Our study represents a significant advancement by providing a comprehensive overview of spatial cell organization in the BM microenvironment of both healthy and diseased states, across mouse models and human subjects. Our work enhances the understanding of the geographic and transcriptional heterogeneity of malignant PC and their associated microenvironment (Fig. 6).

**Fig. 6.**
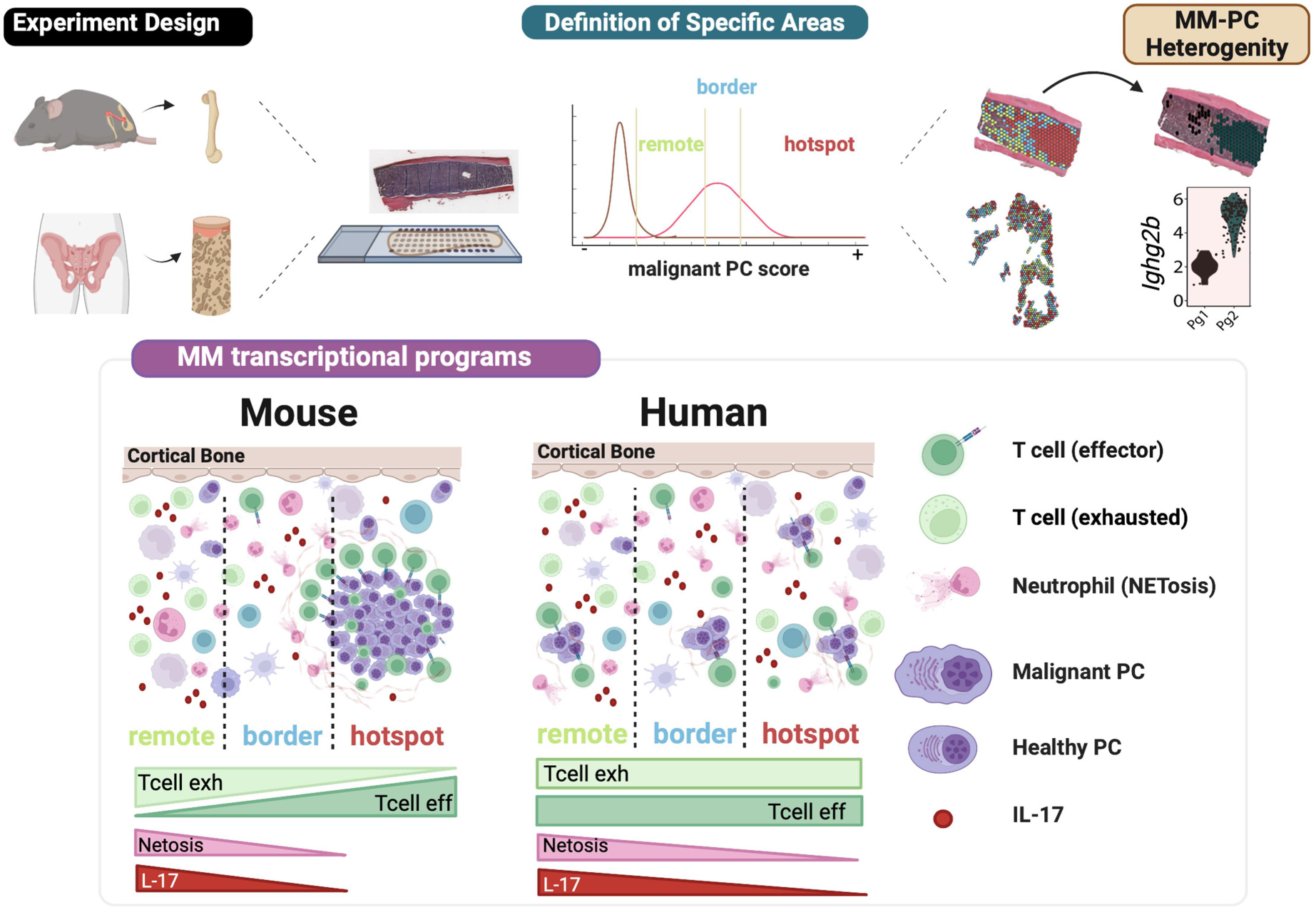
Summary Figure.

The application of spatial technologies is endowed with significant technical challenges. Our initial efforts were focused on preserving RNA integrity during decalcification and using FFPE tissues, first optimized in mouse femurs and then in human samples. As current spatial technologies not always have a single-cell resolution, the use of scRNA-seq data for deconvolution analysis was a requirement in our work using the 10X Visium CytAssyst platform. Other technologies based on targeted sequencing may provide single-cell or even sub-cellular resolution but have other limitations, such as not providing a genome-wide transcriptome analysis, allowing the profiling of significantly smaller regions, or inability to generate quality profiles in calcified tissues, among others.

Using spatial transcriptomics technology on healthy mouse femur tissues, we revealed novel insights into the spatial organization of different BM cell types. These findings align with previously known properties, such as the dominant role of neutrophils near blood vessels in healthy samples^42^ and the correlations between different cell types pointing toward different surrounding areas within the BM microenvironment. The use of malignant PC canonical markers allowed us to identify distinct areas based on the proportion of MM-PC defined as Hotspot, Border, and Remote zones. We acknowledge that in some cases, the expected concentric organization of zones (Hotspot surrounded by Border and Border surrounded by Remote zones) was not observed (e.g., in samples MM1 and MM4). This may reflect the technical limitations of the technology and the heterogeneous nature of the disease. Additionally, we characterized the spatially heterogeneous distribution of transcriptionally diverse malignant PC groups within areas of high tumor cell density. Interestingly, the observed differences in gene expression did not correlate with disease stage, as demonstrated in the MM1 and MM3 samples. Despite similar percentages of MM-PC infiltration, as determined by flow cytometry (Table S1), the malignant PC populations exhibited distinct expression profiles of genes implicated in MM development and prognosis, including B2m, Ly6k and Cyba^43^. These variations may represent a critical factor underlying the heterogeneity in disease behavior among patients. However, a more detailed and targeted analysis is necessary to elucidate the underlying mechanisms responsible for these differences.

These results are aligned with previous single-cell technologies, demonstrating the significant genetic and genomic heterogeneity between and within tumors^44,45^. An advantage of spatial transcriptomics is the possibility to add spatial information to changes in transcriptional programs. As shown in our study, we characterized transcriptional programs involved in the pathogenesis of MM within the BM microenvironment that were associated with MM-PC density gradient. The spatial location of T effector cells was preferably around Hotspot areas, while the T exhausted phenotype was enriched in Remote and Border zones. We also identified the upregulation of several pathways in the malignant PC microenvironment indicative of its transcriptional remodeling in MM disease. The implications of IL-17 signaling, atherosclerosis, osteoclast differentiation, and the involvement of neutrophils through the NETosis mechanism contribute to the malignant PC microenvironment’s spatial heterogeneity.

Our study was further validated in human samples by applying spatial transcriptomics to FFPE-BM biopsies from healthy and MM patients with various degrees of MM-PC infiltration. This validation underscores the reliability and relevance of our murine model’s findings. Like the findings in diseased mouse samples, we identified three distinct regions based on the spatial expression of malignant PC. The potential compromised structural integrity of human archive samples along with the fact that trephine biopsies only represent a limited section of the bone, unlike mouse bone samples, may have an impact on the delineation of the different areas of MM-PC. This technical limitation may explain the differences in T cell behavior between humans and mice. In spite of this, we cannot rule out that differences in disease stage (human samples were obtained from patients at diagnoses and relapse) may also be responsible for these differences. On the other hand, we observed similar transcriptional changes in the diseased BM between mice and humans, particularly in pathways related to NETosis and IL-17 signaling, among others. This consistency was evident as we transitioned from the Remote zone to the Hotspot region in human subjects, providing insights into how the spatial organization influences the genetic activity involved in MM development and progression^8,9,46–48^.

While our investigation offered valuable observations of myeloma’s spatial and transcriptional heterogeneity, we acknowledged the study’s limitations. Firstly, a small number of samples were used. This limited the mechanistic insights, highlighting the importance of increasing sample sizes for more robust conclusions. The low resolution and dependence on single-cell data also emphasizes the need for future experiments using higher-definition resolution technologies. Furthermore, while we examined tissues in three dimensions, our study only assessed one slide per sample, potentially missing critical spatial variations within the tissue. MM is characterized by clonal immunoglobulin V(D)J signatures, originated during the normal process of B cell development, and driving an oncogenic immunoglobulin phenotype commonly present in premalignant conditions in similar percentages^49^. However, the current technology cannot conduct specialized analyses such as V(D)J recombination. Lastly, the importance of studying all the MM-PC microenvironment components remains, even though non-hematopoietic components represent less than 1% of the total BM. Addressing these limitations in future research will be crucial for advancing our understanding of myeloma pathogenesis and facilitating therapeutic development.

In conclusion, this research significantly contributes to the biology of MM within the complex BM tissue. It introduces a novel approach for identifying spatial interactions between MM-PC and their BM microenvironment. This advancement could be pivotal for uncovering mechanisms of resistance and evaluating the efficacy of advanced therapies such as Chimeric antigen receptor T (CAR T) cells. Future studies should aim to elucidate the mechanisms underlying the focal bone lytic lesions observed, to better understand their development and impact. Additionally, exploring the role of malignant PC groups in the context of disease relapse is crucial for a more comprehensive understanding of MM progression.

## METHODS

### Mice and human samples

For mouse studies, bone femur samples were obtained from control (YFPcγ1) and MM (MIcγ1)^21^ derived mice females at age 4-12 months and submitted to Spatial transcriptomics interrogation. Animal experimentation was carried out following the European Communities Council Directive (2010/63/UE) and with the approval of the Ethical Committee of Animal Experimentation of the University of Navarra and the Health Department of the Navarra Government. For human studies, archived paraffined BM patient biopsies of six MM patients (47–78 years of age) and three fresh- collected elderly healthy controls were obtained from Pathology Department of Clínica Universidad de Navarra (CUN) and Hospital Universitario de Navarra, respectively, after written informed consent was achieved. The human sample collection and research conducted in this study were approved by the Research Ethics Committee of the University of Navarra. Personal data was kept confidential following the Organic Law 3/2018 on personal data protection and Spanish Law 14/2007 on Biomedical research. All collection samples are codified; only authorized personnel can correlate the patient’s identity with the codes.

### Immunohistochemistry

Tissue sections from BM femurs from mice were stained using an automated immunostaining platform (Discovery XT-ULTRA, Ventana-Roche). Briefly, after the antigen retrieval with Tris-EDTA (pH=9) for 30 min at 95 °C, sections were stained with anti-GFP antibody (NB100-1770S, 1:2000 dilution, Novus Biologicals), and slides were incubated with the rabbit anti-goat Biotinylated (E0466, 1:200; DAKO) and the visualization system Envision anti Rabbit (K4033; DAKO) conjugated to horseradish peroxidase. IHC reactions were developed using 3,3′-diaminobenzidine tetrahydrochloride (K3468; DAKO). Finally, nuclei were counterstained in Harris′ Hematoxylin. The same process was used to reveal the presence of CD138 (B-A38) or CD38 (SP149) positive cells in the human biopsies but with the visualization systems (Omni-Map anti-Rabbit) conjugated to horseradish peroxidase developing the reactions with the 3,3′-diaminobenzidine tetrahydrochloride (ChromoMap DAB, Ventana, Roche) and purple chromogen (Discovery Purple Kit, Ventana, Roche).

### Spatial transcriptomics library preparation and sequencing

Bone femur mice samples were fixed in formol 4% (PanReac) for 24 h at room temperature (RT), and then decalcified with EDTA 0.25 M pH 6.95 (Invitrogen) for 10 days at RT with agitation. After decalcification, samples were washed in distilled water for 5 min and sequentially dehydrated in ethanol 70% (1 h), 80% (1 h), 96% (1 h), and 100% (overnight), followed by 4 h in xylol (Panreac). The femurs were subsequently embedded in paraffin and incubated at 60 °C overnight. Next, 5 µm tissue sections were mounted on microscopy slides, baked at 42 °C for 3 h, and dried in a desiccator at 37 °C overnight. For spatial transcriptomics assays, preparations were deparaffined, rehydrated, and H&E stained following 10X Genomics recommendations. Tissue imaging was performed using an Aperio CS2 Scanner (Leica Biosystems) at 20x magnification. Sections were destained and decrosslinked before library construction using Visium CytAssist Spatial Gene Expression for FFPE Mouse Transcriptome (10X Genomics). Briefly, a whole transcriptome panel consisting of probe pairs was hybridized with their target RNAs in the tissue sections. Neighboring probe pairs that had hybridized to RNA were then ligated. These ligated probes were then released through RNase treatment and tissue permeabilization and transferred to Visium slides containing spatially barcoded oligonucleotides using a CytAssist instrument (10X Genomics). Probes were spatially labeled through extension, released from the slide, and pooled. Next, samples were indexed via PCR amplification. The resulting libraries were quantified with Qubit dsDNA HS Assay Kit, and their profile was examined using Agilent’s HS D1000 ScreenTape Assay. Sequencing was carried out in an Illumina NextSeq2000 using paired-end, dual-index sequencing (Rd1: 28 cycles; i7: 10 cycles; i5: 10 cycles; Rd2: 50 cycles) at a minimum depth of 25000 reads per spot. Per the manufacturer’s instructions, archived human FFPE-BM patients’ biopsies mounted on microscopy slides were subjected to spatial transcriptomic analysis using Visium CytAssist Spatial Gene Expression for FFPE Human Transcriptome (10X Genomics). The protocol workflow resembles the one described above, except tissue sections are directly mounted on Visium slides. Hence, upon the release of the probes that follow RNase treatment and tissue permeabilization, they directly diffuse onto the spatially barcoded oligonucleotides, precluding the need for a CytoAssist instrument. In this case, the whole human transcriptome is covered with 3 probe pairs per targeted RNA.

### Preprocessing and QC spatial transcriptomics data

Spatial transcriptomic data were demultiplexed and mapped using the SpaceRanger^50^ software (v2.0.1). The Ensembl 105 genomes were used as reference (GRCh38 for humans and mm10 for mice samples). Filtered feature-barcode expression matrices obtained from SpaceRanger were used as initial input for the spatial transcriptomics analysis using Seurat^51^ (v4.3.0.1) and STutility^52^ (v1.1.1). Spots overlapping the cortical region of the bone were manually removed based on the H&E staining images usingCloupe (v6.0.0). For mouse samples, SCTransform^53^ (v0.3.5) was used for normalization, using the regularized negative binomial regression, and for regressing out between-sample variations. Afterward, spots with less than 200 Unique Molecular Identifier (UMI) (also called counts), and 150 features (genes) were filtered out. Human samples were normalized with SCTransform^53^ (v0.3.5), and spots corresponding to bone tissue were manually removed based on the H&E images using Cloupe (v6.0.0).

### Single-cell RNA-seq data in mice

We downloaded a preprocessed murine single-cell dataset from https://apps.embl.de/nicheview/, published by Baccin et al^24^, for deconvolution analysis. We used the annotation contained in Baccin et al^24^; we preselected six highly prevalent cell types for the reference (neutrophils, DC, T cells, erythroblast, B cells, and monocytes). Therefore, low prevalent cell types were filtered out or relabeled (e.g., all T cell subtypes into one); as a result, we generated a single cell set for the spot deconvolution of healthy murine BM. For the deconvolution of the spatial mouse MM samples, we included an additional cell type, MM-PC, obtained from Cenzano et al^25^. The murine scRNA-seq dataset^24^ used previously for healthy mice with the new data set from malignant PC^25^ were integrated using Harmony (v1.2.0) with default parameters. We denote that integrated set as scBMReference.

### Deconvolution analysis and hierarchical clustering in mice

Deconvolution analysis was performed in mice spatial transcriptomics data using scBMReference as the reference for CARD^54^ (v1.0.0) with default parameters. The estimated cell-type percentage values derived from the deconvolution were used to cluster spots using the hierarchical clustering implementation in factoextra^55^ (v1.0.7) with default parameters. The optimal number of clusters was determined by evaluating the consistency of marker gene expression profiles and employing the silhouette method; as a result, k=4 and k=7 was defined as the number of clusters in healthy and malignant samples, respectively. Each cluster’s most abundant cell types were calculated by averaging the cell types contributing at least 20% to each spot signature. Correlation analyses were conducted using corrplot^56^ (v0.92). Note that the prevalence analysis in Fig. 2b of MM-PC in healthy samples was conducted as a quality control, but they were not included in the clustering analysis. In dot-plots showing the average of the most abundant cell types and in the pie-chart-based spatial visualization, only cell types contributing at least 20% to each spot based on the deconvolution signature were considered. The proportion of these cell types was recalculated based on this filtering.

### Spot Labelling and Plasma Group Identification in mice

We obtained the top marker genes (Table S2) by comparing the transcriptome of the MM-PC versus the rest of the cell types included in our scBMReference, using the default Seurat pipeline. Then, we filtered out the genes that had no expression in our spatial transcriptomic datasets, both human and mouse samples. Malignant PC score was calculated using AUCell (v1.18.1), which computes the area under the curve for a given gene-set (a unitless measure) per spot. This score, also called signature, reflects the activity of a gene-set, in this case, the MM-PC marker genes. All the spots with a lower PC score than the value equivalent to the 90% percentile obtained in the healthy samples were removed from the next annotation steps and labeled as “Rest” (Fig. S5a). Then, for each mice MM sample independently, the remaining spots were divided based on the malignant PC score using percentiles and labeled as “Remote zone” (malignant PC score < 25% percentile), “Border zone” (25% percentile < malignant PC score < 50% percentile) and “Hotspots” (malignant PC score > 50% percentile) (Fig. 3b). The labeling Hotspot-Border zone-Remote zone, denotes the expected concentric organization of these zones (with Hotspot surrounded by Border zone, and Border zone by Remote zone); however, this was not the case for all the spots in all mice samples. "Hotspot" spots were then grouped based on their spatial position using the igraph package (v1.5.1). Each spot was connected to its six nearest neighbors in the spatial configuration, reflecting their designed spatial distribution within the 10X Visium dataset. The silhouette score was employed to identify the appropriate resolution for modularity (Fig. S5b-c). As a result, “Hotspot” spots were grouped into twelve PC groups (Pg) (Fig. 3c).

### Plasma Group Variability analysis in mice

Gene expression variability analysis within Pg was conducted using pseudo-bulk profiles. Reads from all the spots within each Pg were added; secondly, the resulting pseudo-count matrix was normalized using DESeq2 (v1.36.0). Standard error (SE) was calculated for each gene across all the Pg, and then genes were ranked based on this SE. The top 1% of genes were labeled highly variable (Fig. S5e). Over-representation analysis (ORA) using GO was performed on the highly variable genes to identify the pathways associated with them using clusterProfiler (v4.4.4) with default parameters.

### Differential analysis between regions

Differential analysis between the three areas (“Remote zone”, “Border zone”, and “Hotspot”) was conducted with the FindMarkers function with default parameters from Seurat (v5.0.1) on the SCT Assay after merging data from the four MM mice samples. Genes were defined as significant if their adjusted p-values were < 0.05. Subsequently, ORA was carried out on genes that were either up- (average log2 fold change > 0) or down-regulated (average log2 fold change < 0) for each of the comparisons (“Remote zone vs Border zone” and “Border zone vs Hotspot”) using the Kyoto Encyclopedia of Genes and Genomes (KEGG) database. Moreover, the area under the curve was computed for several pathways using the AUCell (v1.18.1) using different gene-sets of interest (Table S2 and Supplementary Data 2 and Supplementary Data 4). Statistical comparisons of pathway scores across different zones were performed using the Wilcoxon statistics.

### Spot Labelling in Human Samples

As previously explained, spots were labeled based on each sample’s malignant PC score within the MM-BM. Given the large variability in UMI counts between and within human samples, a preprocessing step was necessary. First, we divided each sample into bins based on the UMI counts, with 11 bins ranging from 0-500 to over 5000 UMIs (Fig. S9a). Within each bin, we applied the per-sample analysis explained for mice (See Spot Labelling in mice). Briefly, spots with a malignant PC score below the 90% malignant PC score obtained from healthy human samples were labeled “Rest” and removed from the next annotation steps. Then, the remaining spots were further classified as “Remote zone” (malignant PC score < percentile 33), “Border zone” (percentile 33 < malignant PC score < percentile 66) or “Hotspot” (malignant PC score > percentile 66) (Fig. S9b). Finally, the new labeling was assigned to the original Seurat Objects, ensuring the integrity of the spatial information. Even though in human samples there is no concentric annotation of these areas, we kept the labeling used in mice for consistency. Pathways scores were computed using the AUCell (v1.18.1) on raw counts for each sample (Table S4 and Supplementary Data 5). We corrected the scores using housekeeping genes (HSIAO_housekeeping_genes) to mitigate potential biases arising from technical differences. Statistical comparisons of pathway scores across different zones were conducted using Wilcoxon statistics, enabling robust assessments of pathway activity.

### Data and code availability

Mouse and human spatial raw and processed data have been deposited at GSE269875. All data will be publicly available as of the date of peer-reviewed publication. Mouse scRNA-seq data used for deconvolution was obtained from Baccin et al^24^ (https://apps.embl.de/nicheview/). All original code has been deposited at https://github.com/lsudupe/spatialMM.

## ACKNOWLEDGEMENTS

This work was supported by the Instituto de Salud Carlos III and co-financed by ERDF A way of making Europe (PI20/01308, PI23/00516); CIBERONC (CB16/12/00489), RICORS TERAV (RD21/0017/0009); Departamento de Industria Gobierno de Navarra (AGATA 0011-1411-2020- 000010/0011-1411-2020-000011) and Departamento de Salud Gobierno de Navarra; the Cancer Research UK [C355/A26819]; FC AECC and AIRC under the Accelerator Award Program; The International Myeloma Foundation (Brian van Novis) and The Paula and Rodger Riney Foundation to FP. The study was also supported by The Spanish Government through project PID2019-111192GA- I00 (MICINN) to DGC. IAC was supported by a Marie Curie grant (H2020- MSCA-IF-837491) from the European Commission. LS was supported by KAUST Discovery Doctoral Fellowship.

We would like to thank the staff of the advanced genomic lab and the animal facility at CIMA Universidad de Navarra for their invaluable technical and intellectual assistance. We also acknowledge the Hospital Universitario de Navarra and the Biobank of the University of Navarra. We are particularly grateful to the patients and healthy donors who participated in this study.

## AUTHORSHIP CONTRIBUTIONS

L.S. and A.R.L.P. analyzed the spatial transcriptomic data. L.S., E.M.L., A.R.L.P., and I.A.C. interpreted the spatial transcriptomic data. E.M.L. achieved all the final figures. L.S. and I.A.C. wrote the paper; A.V., S.S., P.S.M., and P.A.R. performed the spatial transcriptomic experiments. P.R.C. performed the histologic analysis. M.L. and J.A.M.C. executed and ceded the animal model for this study. J.R. and I.S.G. provided the BM samples from healthy adult donors. L.A.G. and M.A. performed the histologic analysis and screening of the human samples. D.G.C., with help from V.L. and J.T., supervised the computational work. E.M.L. conceived the research project. I.A.C., D.G.C., and. F.P. conceived and directed the research project. All authors actively participated in the discussions underlying this manuscript. L.S., E.M.L., A.R.L.P., I.A.C., D.G.C., and F.P. discussed the results and edited the final manuscript. All authors contributed to reading and approving the final manuscript.

## SUPPLEMENTARY FIGURES

**Supplementary Figure 1.**
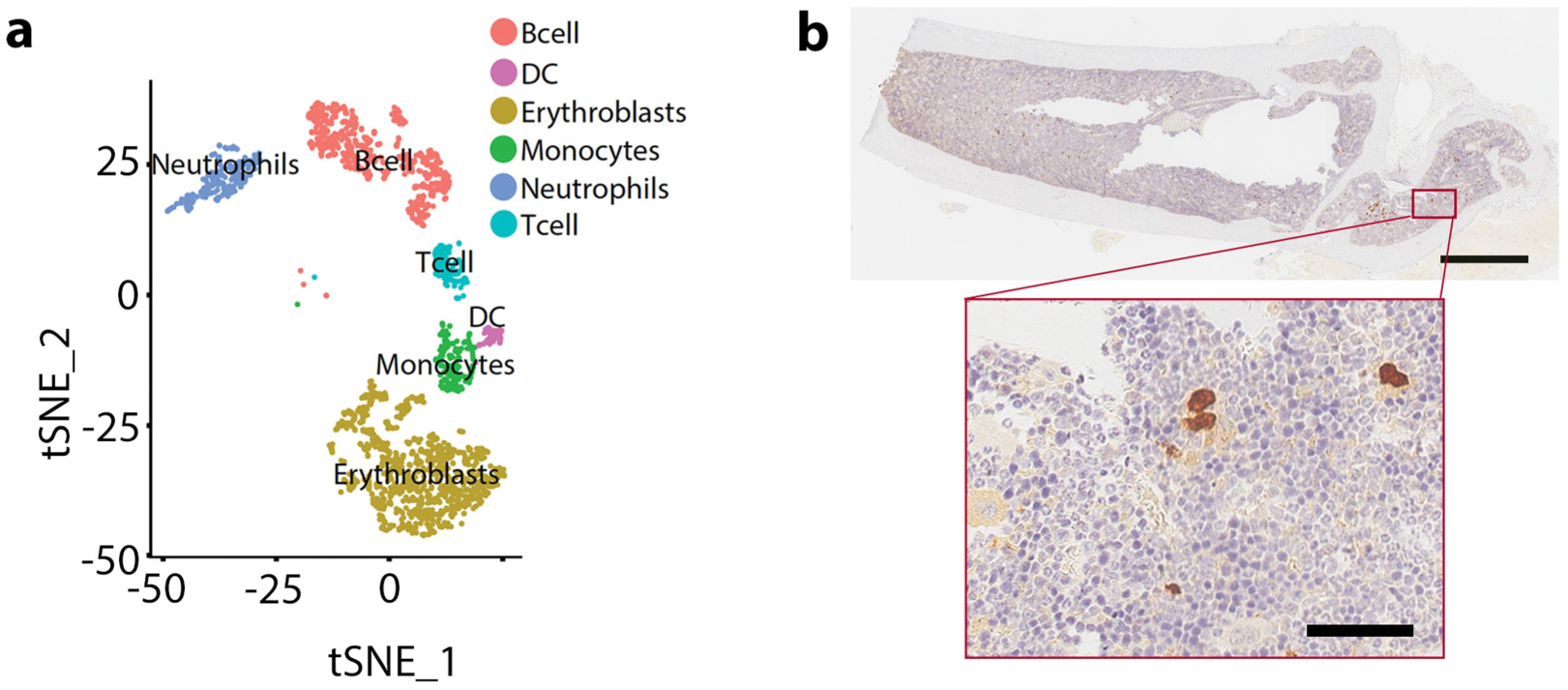
Spatial characterization of healthy bone tissue. **(a)** UMAP representation of the main identified cell types by deconvolution analysis. **(b)** H&E staining combined with a GFP immunostaining for healthy PC visualized with DAB. Scale 1000 μm (upper) and 100 μm (bottom).

**Supplementary Figure 2.**
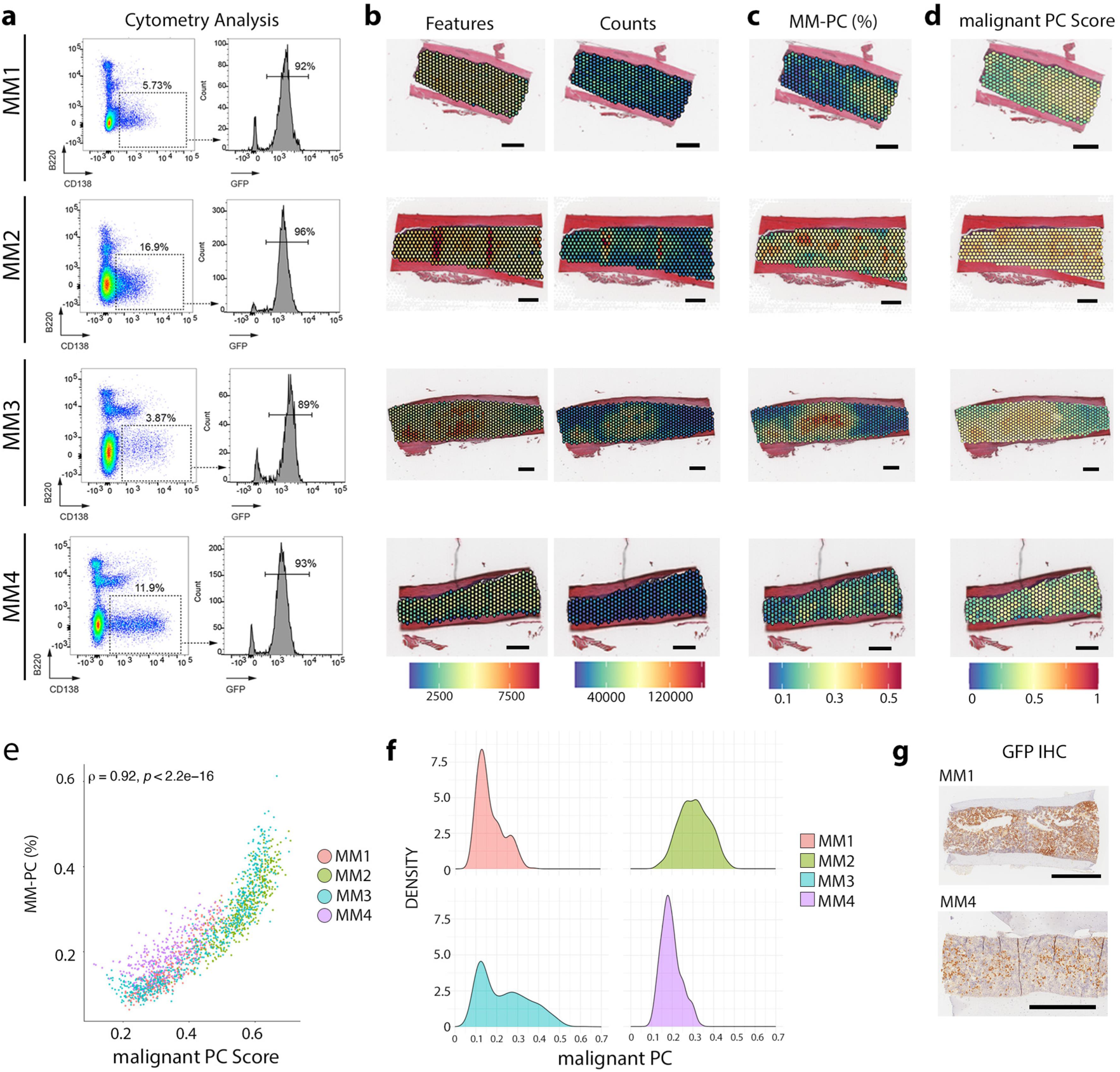
Additional information of the malignant bone marrow samples. **(a)** Cytometry analysis of MM-PC infiltration percentage through the GFP^+^CD138^+^B220^-^, **(b)** quality control of the diseased bone tissues, Features, and Counts respectively, **(c)** anatomical position of MM-PC, and **(d)** spatial distribution of the malignant PC enrichment score in the diseased samples (MM1, MM2, MM3, and MM4). **(e)** Correlation between the MM-PC and the malignant PC enrichment score. **(f)** Malignant PC signal density plots of diseased femur sections. **(g)** GFP IHC of the malignant samples (MM1 and MM4). Scale of 500 μm (b-d) and 1mm (g).

**Supplementary Figure 3.**
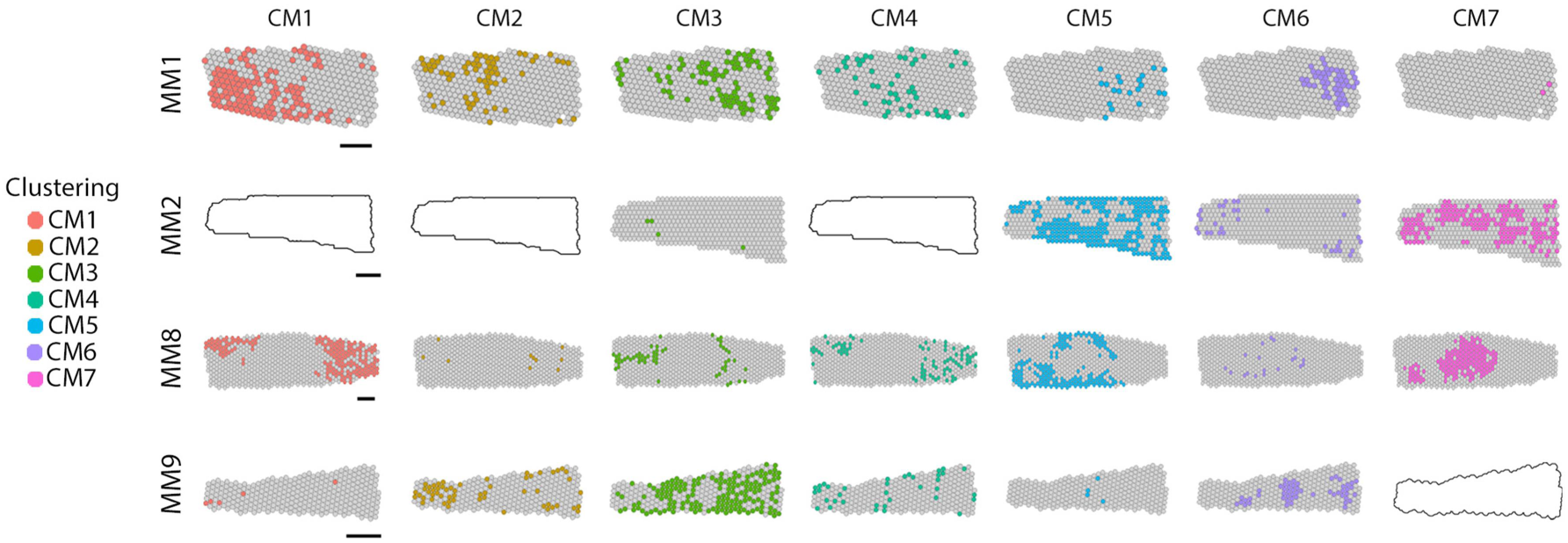
Spatial characterization of diseased samples. Spatial individualized visualization of the seven identified clusters (CM1-CM7) in the pathologic bone femur sections (MM1, MM2, MM3, and MM4). Scales of 500 μm.

**Supplementary Figure 4.**
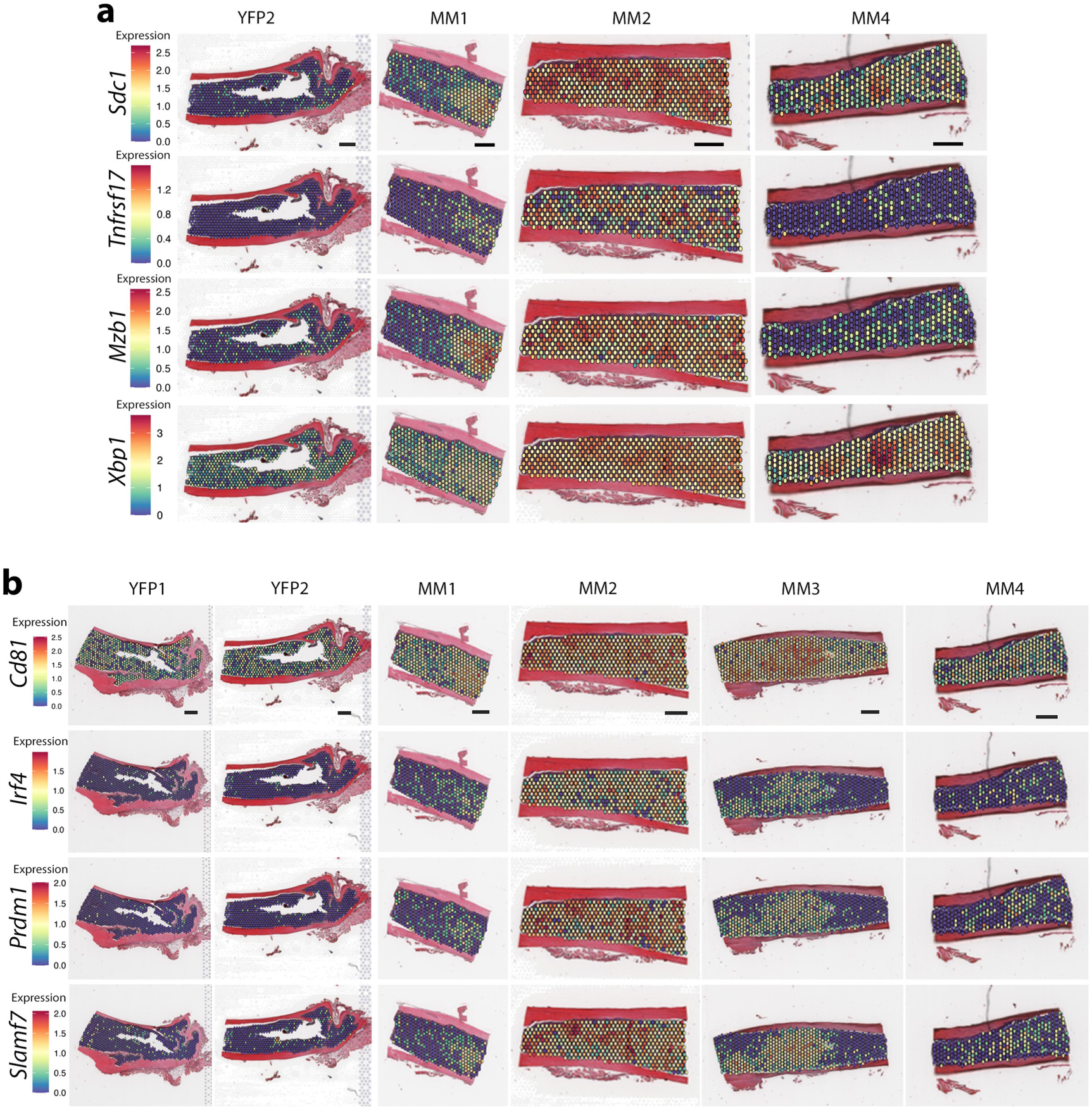
Spatial expression of pathologic plasma cell canonical markers. **(a)** Additional information to the main figure in YFP2, MM1, MM2, and MM4 samples and **(b)** selected pathological markers in all the samples, both healthy (YFP1, YFP2) and pathologic (MM1, MM2, MM3, and MM4) samples. Scales 500 μm.

**Supplementary Figure 5.**
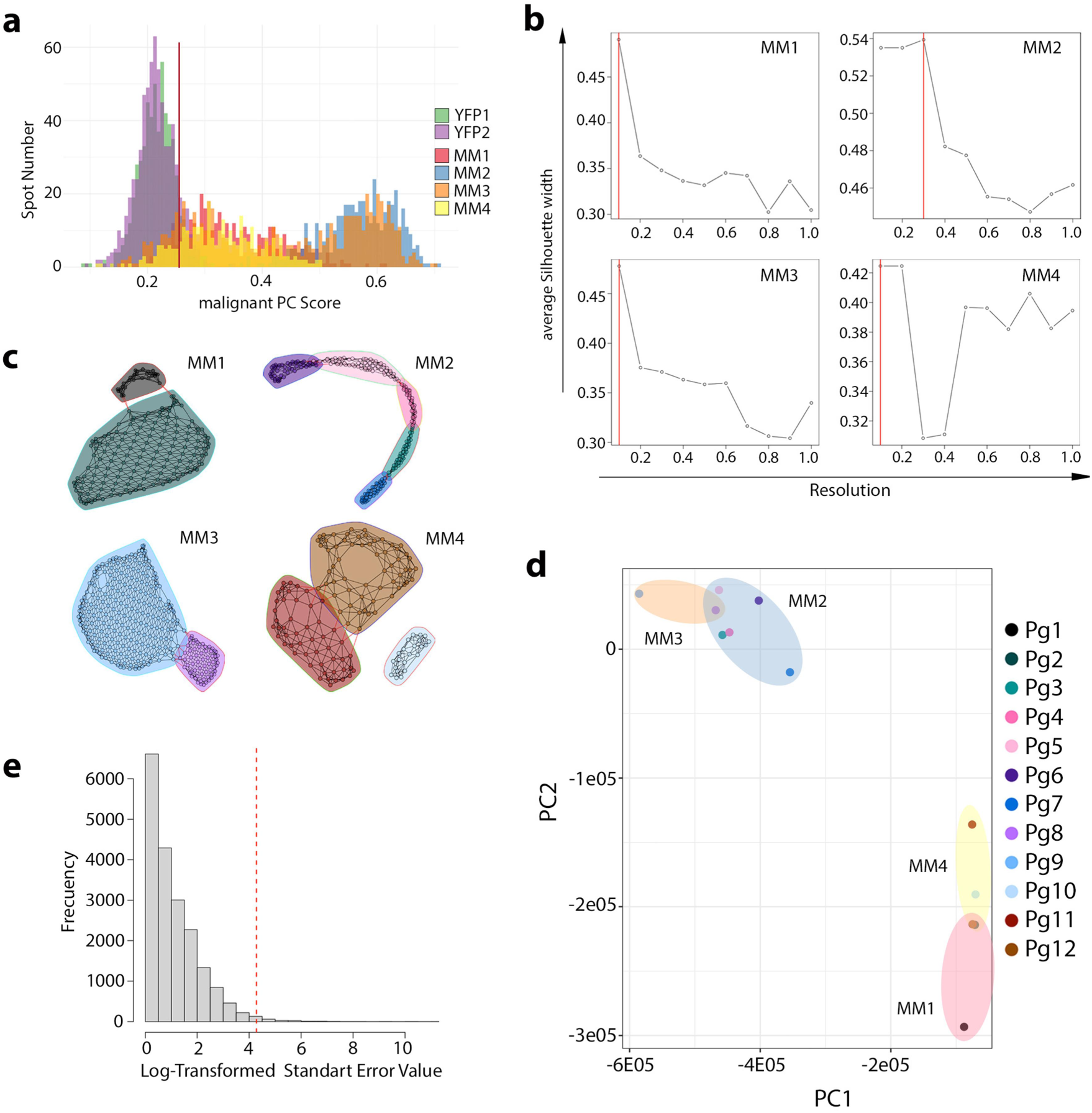
Definition of the diseased samples’ different areas and plasma cell groups. **(a)** Histogram of malignant PC enrichment score distribution in all the samples. **(b)** Silhouette width score for modularity. **(c)** Spatial graph subclusters of hotspots from malignant samples. **(d)** Principal component analysis (PCA) of the twelve identified MM-PC groups (Pg1-Pg12) in the diseased samples. **(e)** Histogram of the standard error (SE) distribution to select the 1% highly variable genes across the MM-Pg groups.

**Supplementary Figure 6.**
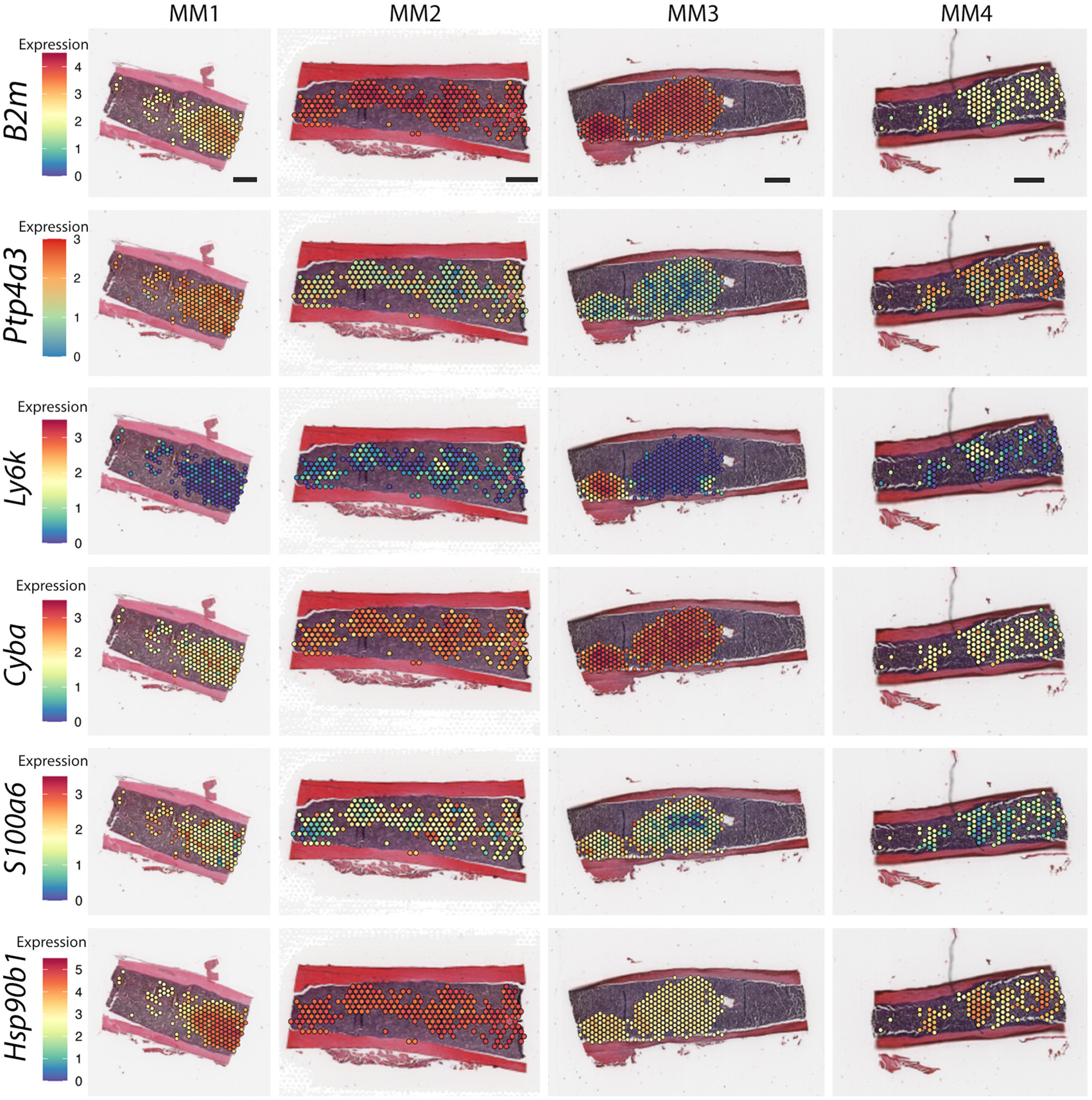
Spatial expression of the selected highly variable genes across MM-PC groups. MM1, MM2, MM3 and MM4. Scales 500 μm.

**Supplementary Figure 7.**
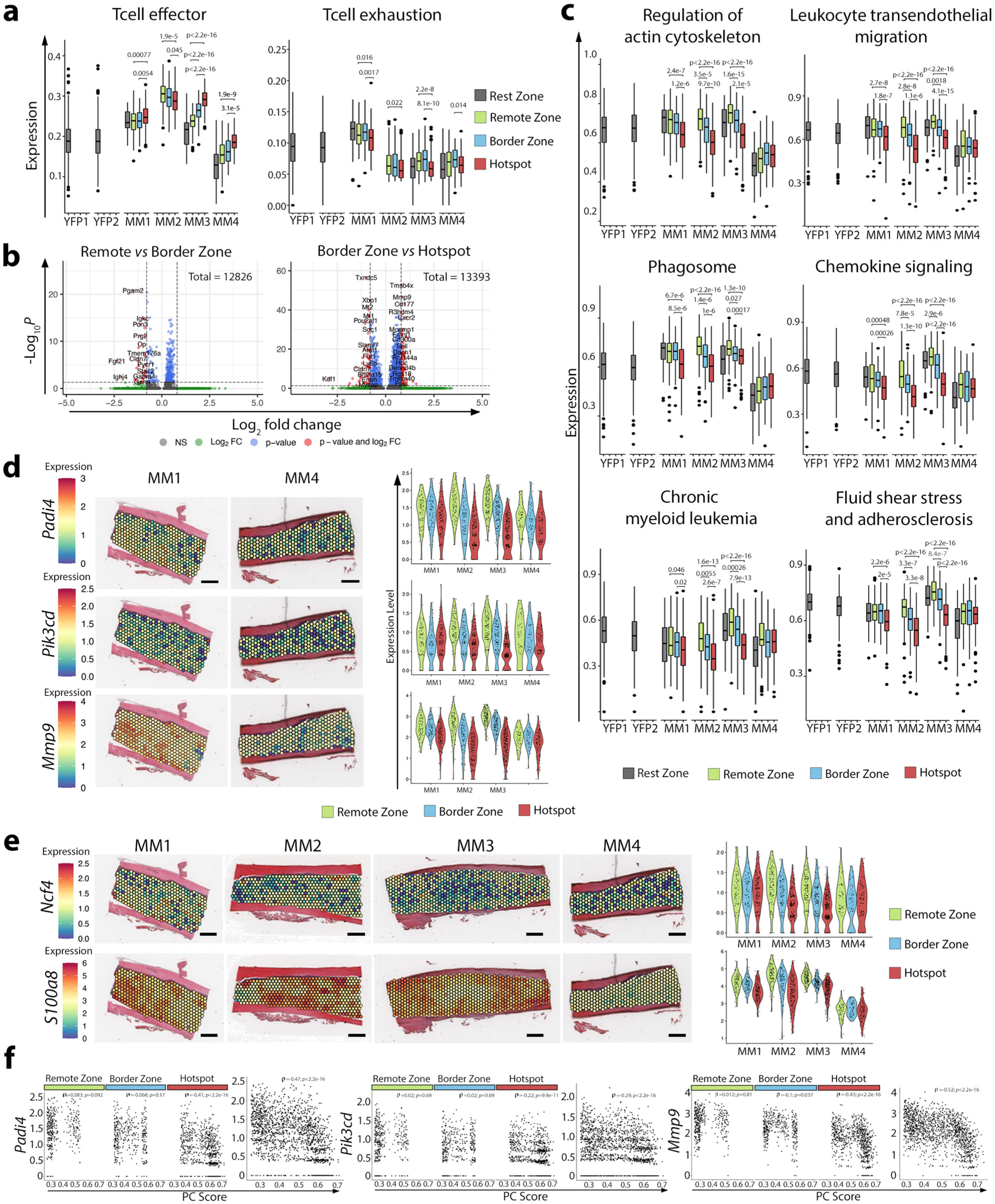
Additional information on the spatial complexity of the MM-PC niche. **(a)** Box plots of the Tcell effector and Tcell exhaustion profiles in all healthy and pathologic samples **(b)** Volcano plots showing DEG between all the identified areas. **(c)** Box plots of the selected KEGG pathways split by sample. **(d)** Spatial distribution of *Padi4*, *Pik3cd*, and *Mmp9* genes in MM1 and MM4 samples. **(e)** Spatial distribution and violin plots of *Ncf4* and *S100a8* genes in all the diseased samples. **(f)** Correlation of the *Padi4*, *Pik3cd*, and *Mmp9* genes with the malignant PC enrichment score. Scales of the spatial gene representations, 500 μm.

**Supplementary Figure 8.**
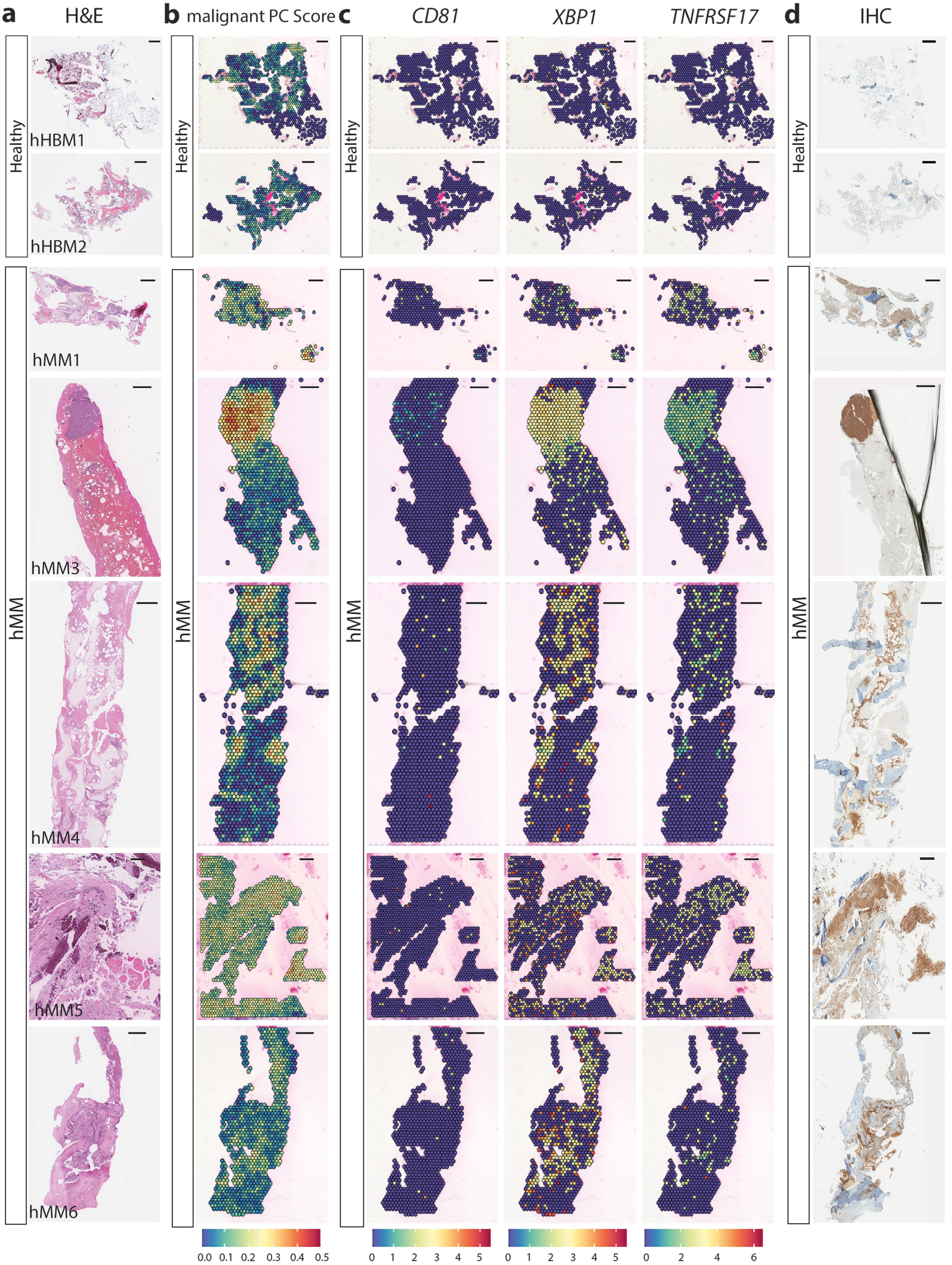
Validation of the spatial technology in human samples. **(a)** H&E staining, **(b)** spatial distribution of the malignant PC enrichment score, **(c)** spatial expression of human malignant PC canonical markers, and **(d)** CD138 (hHBM1, hHBM2, hMM1, hMM3, hMM4) and CD38 (hMM2, hMM5, and hMM6) IHC of healthy (hHBM) and diseased human (hMM) bone biopsies. Scales 500 μm.

**Supplementary Figure 9.**
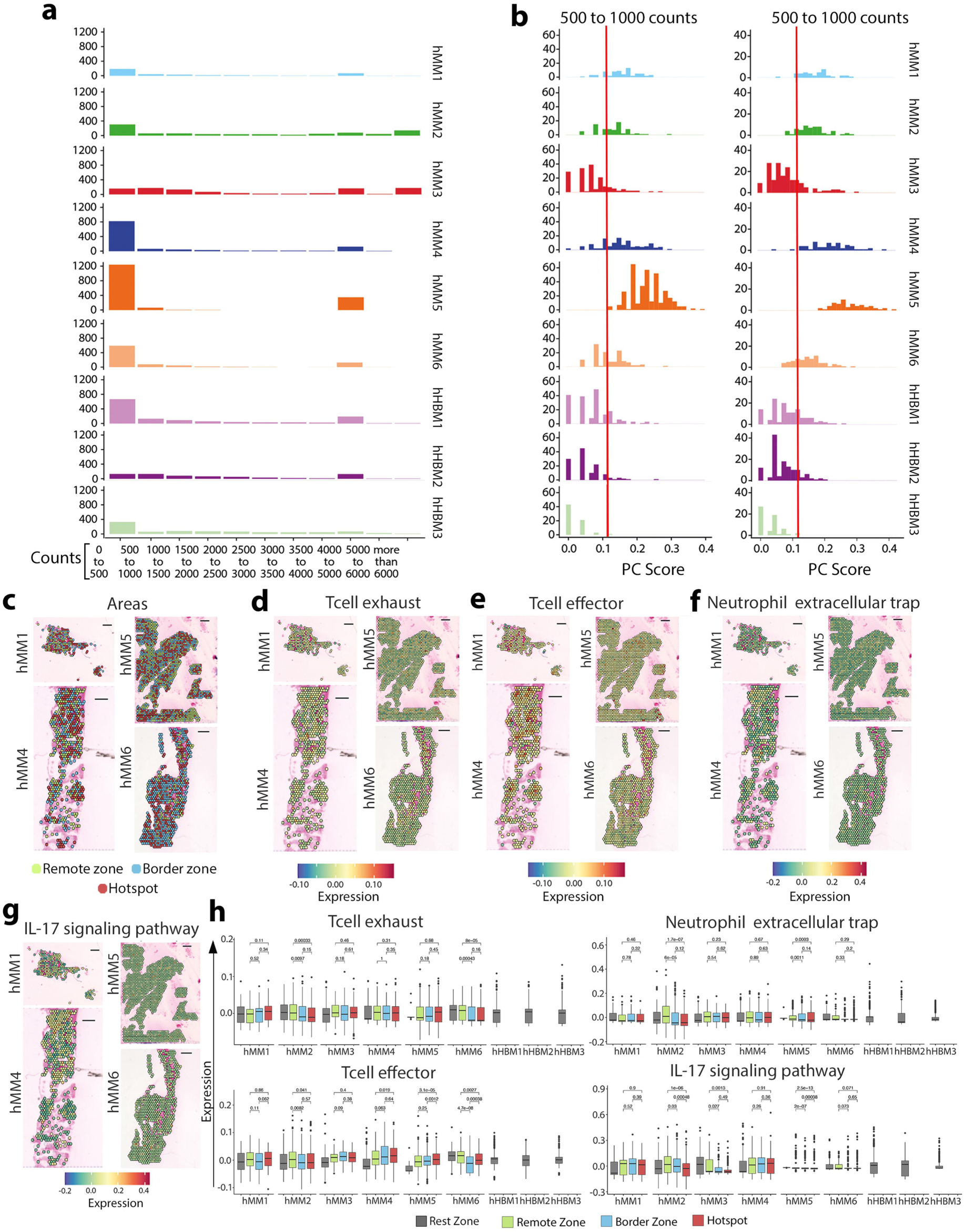
Additional information to Figure 5. **(a, b)** Analytical pipeline for area definition. **(c)** Projection of the three identified areas (Remote zone, Border zone, and Hotspot) in all the human MM samples. **(d-g)** Spatial distribution of the T cell exhaustion (d) and T cell effector scores (e), Neutrophil extracellular trap formation (NETs) (f) and IL-17 signaling pathways (g) in the diseased human samples. **(h)** Box plots of the mentioned profiles split by sample. Scales 500 μm.

**Supplementary Figure 10:**
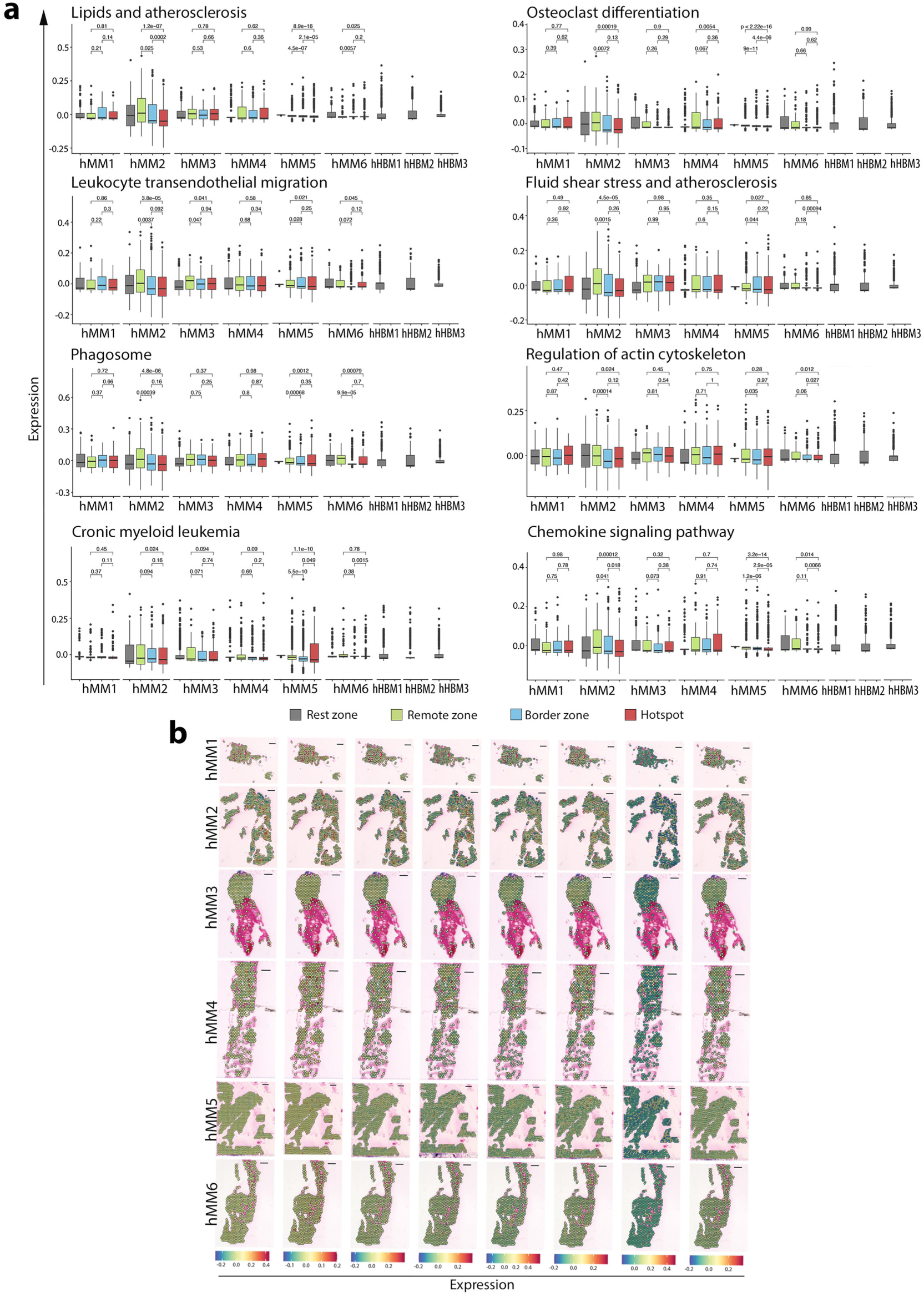
Additional information to figure 5. (a) Box plots of the upregulated KEGG pathways represented independently in all human pathologic (hMM1-hMM6) and healthy samples. **(b)** Spatial distribution of the upregulated KEGG pathways represented in all human pathologic samples. Scales 500 μm.

## SUPPLEMENTARY TABLES

**Table S1:** Information of the healthy and malignant mouse femur tissues included in the study.

| **Sample Identification** | **Sex** | **Disease** | **Tumoral cells<sup>1</sup>**<br>**(%)** | **DV 200<sup>2</sup>**<br>**(%)** |
| --- | --- | --- | --- | --- |
| YFP1 and 2 | Female | Healthy | 0.23 | 82.55 |
| MM1 | Female | MM | 5.3 | 73.78 |
| MM2 | Female | MM | 16.22 | 69 |
| MM3 | Female | MM | 3.44 | 45 |
| MM4 | Female | MM | 11.01 | 74 |

<sup>1</sup>Tumoral cells: Percentage of GFP and CD138 positive cells and B220 negative cells.
<sup>2</sup>DV200: Percentage of total RNA fragments > 200 nucleotides

**Table S2:**
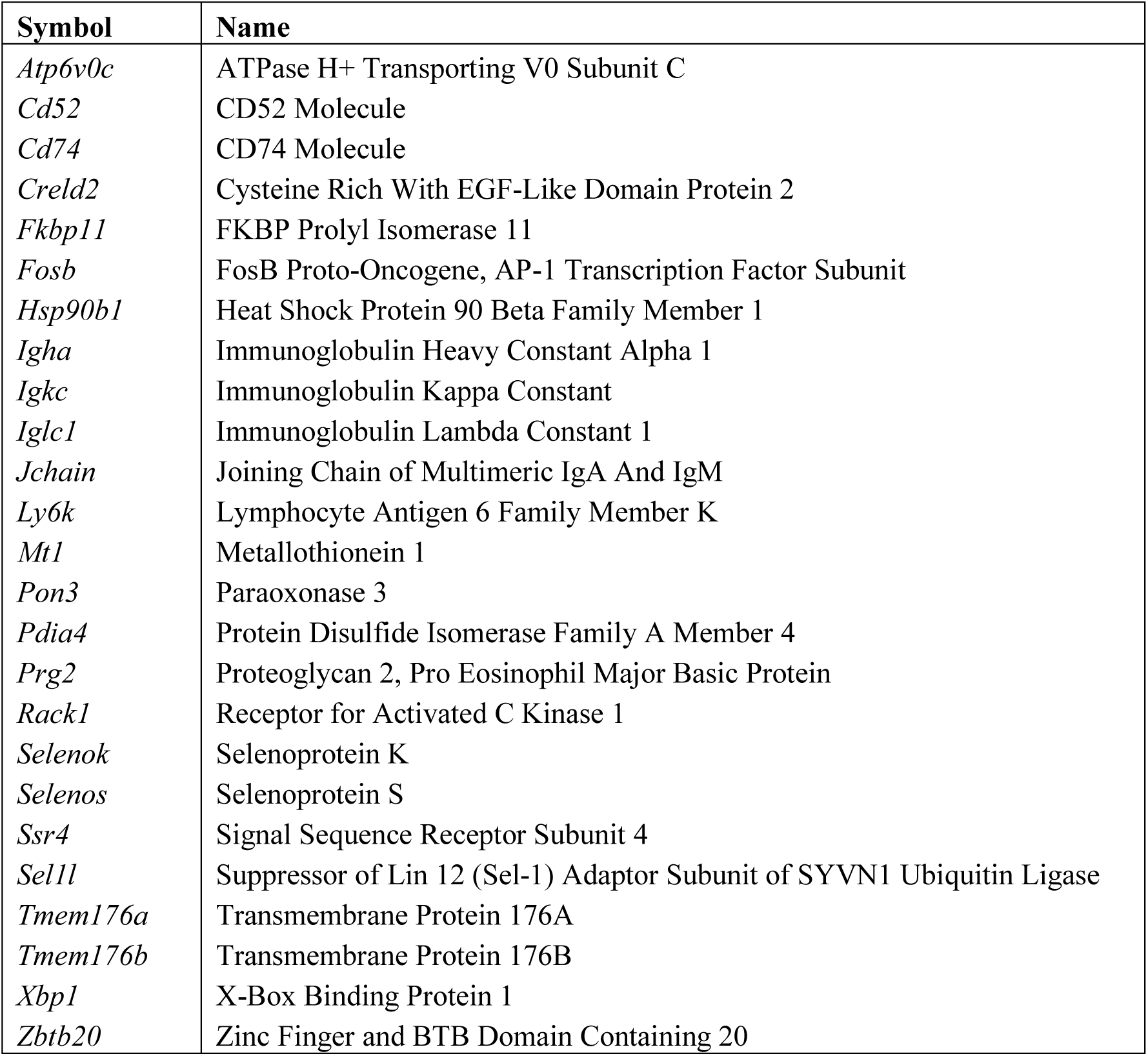
Selected genes for calculating the malignant plasma cell enrichment score in mouse.

**Table S3:**
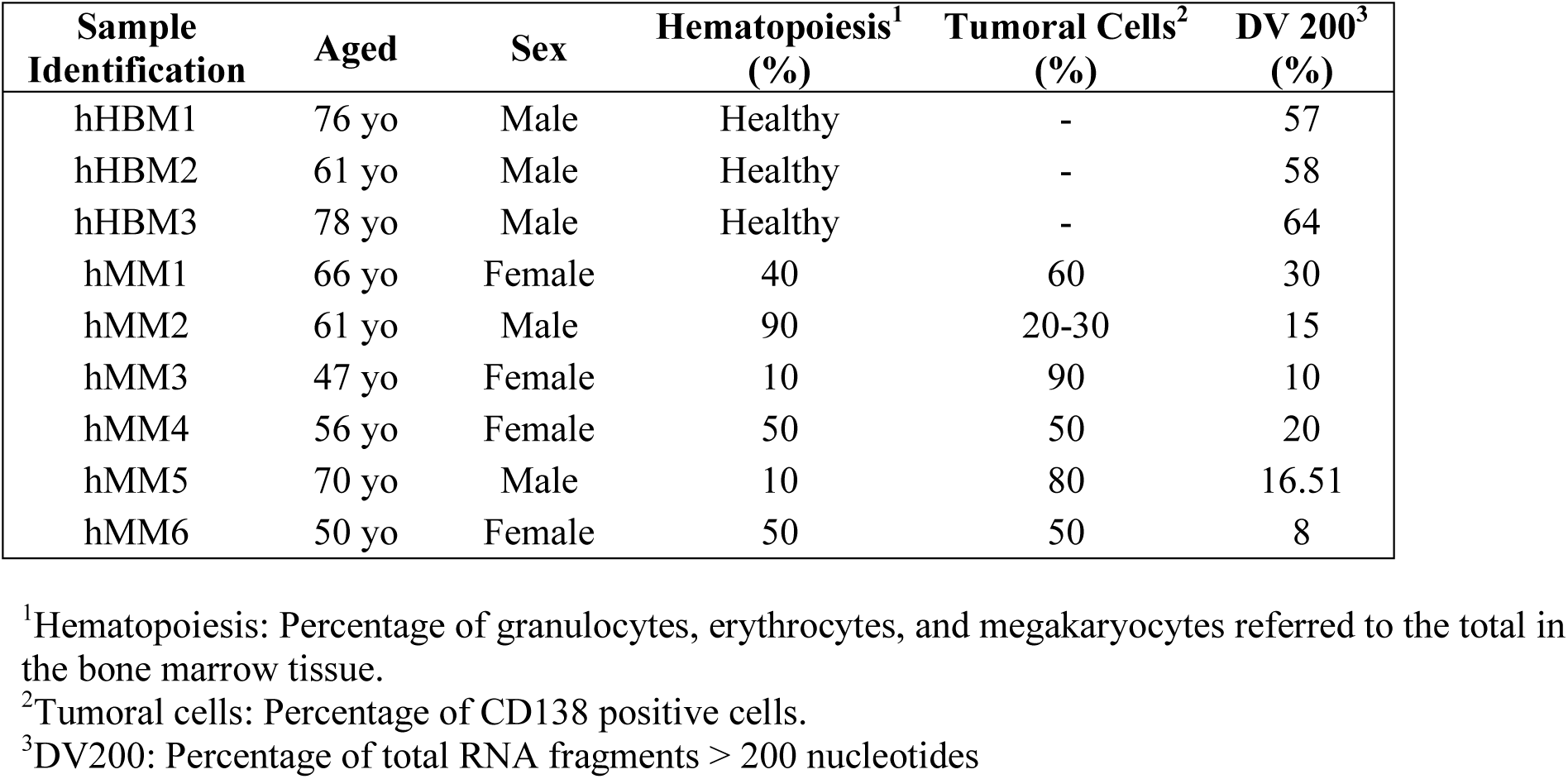
Information of the healthy and malignant human bone tissues included in the study.

**Table S4:**
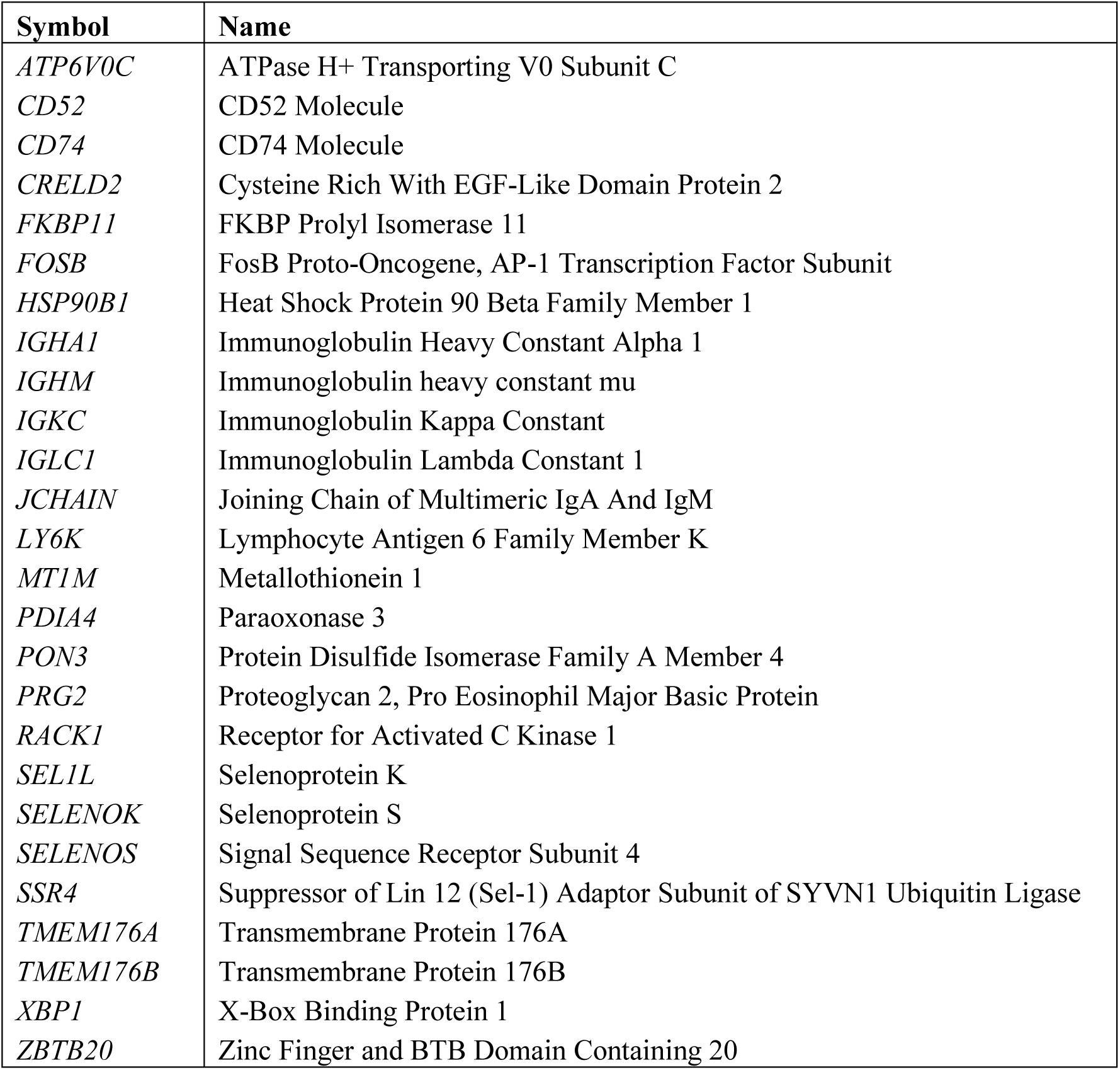
Gene-set used for calculating the malignant plasma cell enrichment score in human.

## SUPPLEMENTARY DATA

**Supplementary Data 1:** Identification of 194 highly variable genes across PC groups (Pg) ranked by standard error (SE).

**Supplementary Data 2:** Gene ontology (GO) terms derived from Over-Representation Analysis (ORA) of the highly variable genes identified across Pg.

**Supplementary Data 3:** Gene-sets used for calculating the T cell exhaustion and T cell effector scores in both mouse and human samples.

**Supplementary Data 4:** Differentially expressed analysis and ORA between the three defined areas, hotspot, border zone, and remote zone.

**Supplementary Data 5:** Gene-sets used for calculating the scores of the enriched pathways in the three defined areas, hotspot, border zone, and remote zone in both mouse and human samples.

## REFERENCES

1. Fröbel, J. et al. The Hematopoietic Bone Marrow Niche Ecosystem. Front. Cell Dev. Biol. 9, (2021).

2. Kumar, S. K., et al. Multiple myeloma. Nat Rev Dis Primers 3, 17046 (2017).

3. García-Ortiz, A. et al. The Role of Tumor Microenvironment in Multiple Myeloma Development and Progression. Cancers 13, 217 (2021).

4. Chen, M. et al. Dynamic single-cell RNA-seq analysis reveals distinct tumor program associated with microenvironmental remodeling and drug sensitivity in multiple myeloma. Cell & Bioscience 13, 19 (2023).

5. de Jong, M. M. E. et al. The multiple myeloma microenvironment is defined by an inflammatory stromal cell landscape. Nat Immunol 22, 769–780 (2021).

6. de Jong, M. M. E. et al. An IL-1β-driven neutrophil–stromal cell axis fosters a BAFF-rich protumor microenvironment in individuals with multiple myeloma. Nat Immunol 1–14 (2024).

7. Binder, A. F., Walker, C. J. & Baljevic, M. Impacting T-cell fitness in multiple myeloma: potential roles for selinexor and XPO1 inhibitors. Front. Immunol. 14, (2023).

8. Van Valckenborgh, E. et al. Multifunctional Role of Matrix Metalloproteinases in Multiple Myeloma: A Study in the 5T2MM Mouse Model. The American Journal of Pathology 165, 869– 878 (2004).

9. Zhao, Z. et al. A neutrophil extracellular trap-related risk score predicts prognosis and characterizes the tumor microenvironment in multiple myeloma. Sci Rep 14, 2264 (2024).

10. Garcia-Gomez, A. et al. Targeting aberrant DNA methylation in mesenchymal stromal cells as a treatment for myeloma bone disease. Nat Commun 12, 421 (2021).

11. Schinke, C. & Weinhold, N. The Immune Microenvironment in Multiple Myeloma Progression at a Single-cell Level. Hemasphere 7, e894 (2023).

12. Quach, H. et al. Early human fetal lung atlas reveals the temporal dynamics of epithelial cell plasticity. Nat Commun 15, 5898 (2024).

13. Fortelny, N. et al. JAK-STAT signaling maintains homeostasis in T cells and macrophages. Nat Immunol 25, 847–859 (2024).

14. Roehrig, et al. Single-cell multiomics reveals the interplay of clonal evolution and cellular plasticity in hepatoblastoma. Nature Communications. 15, 3031 (2024).

15. Singh, V. M. et al. Analysis of the effect of various decalcification agents on the quantity and quality of nucleic acid (DNA and RNA) recovered from bone biopsies. Annals of Diagnostic Pathology 17, 322–326 (2013).

16. Xiao, X. et al. Spatial transcriptomic interrogation of the murine bone marrow signaling landscape. Bone Res 11, 1–13 (2023).

17. Bandyopadhyay, S. et al. Mapping the cellular biogeography of human bone marrow niches using single-cell transcriptomics and proteomic imaging. Cell S0092–8674(24)00408–2 (2024).

18. Tower, R. J. et al. Spatial transcriptomics reveals a role for sensory nerves in preserving cranial suture patency through modulation of BMP/TGF-β signaling. Proc Natl Acad Sci U S A 118, e2103087118 (2021).

19. Tower, R. J. et al. Spatial transcriptomics reveals metabolic changes underly age-dependent declines in digit regeneration. Elife 11, e71542 (2022).

20. John, M. et al. Spatial transcriptomics reveals profound subclonal heterogeneity and T-cell dysfunction in extramedullary myeloma. Blood, 2024024590 (2024).

21. Larrayoz, M. et al. Preclinical models for prediction of immunotherapy outcomes and immune evasion mechanisms in genetically heterogeneous multiple myeloma. Nat Med 29, 632–645 (2023).

22. Ståhl, P. L. et al. Visualization and analysis of gene expression in tissue sections by spatial transcriptomics. Science 353, 78–82 (2016).

23. Hensel, J. A., Khattar, V., Ashton, R. & Ponnazhagan, S. Characterization of immune cell subtypes in three commonly used mouse strains reveals gender and strain-specific variations. Lab Invest 99, 93–106 (2019).

24. Baccin, C. et al. Combined Single-Cell and Spatial Transcriptomics Reveals the Molecular, Cellular and Spatial Bone Marrow Niche Organization. Nature cell biology 22.1 (2020)

25. Cenzano, I. et al. Transcriptional Characterization of the Stromal and Endothelial Bone Marrow Microenvironment during Progression from MGUS to Multiple Myeloma. Preprint at 10.1101/2024.04.24.589777 (2024).

26. Schmidt-Hieber, M. et al. CD117 expression in gammopathies is associated with an altered maturation of the myeloid and lymphoid hematopoietic cell compartments and favorable disease features. Haematologica 96, 328–332 (2011).

27. Bouchnita, A., Eymard, N., Moyo, T. K., Koury, M. J. & Volpert, V. Bone marrow infiltration by multiple myeloma causes anemia by reversible disruption of erythropoiesis. Am J Hematol 91, 371–378 (2016).

28. Leone, P. et al. Dendritic cells accumulate in the bone marrow of myeloma patients where they protect tumor plasma cells from CD8+ T-cell killing. Blood 126, 1443–1451 (2015).

29. Rasche, L. et al. Spatial genomic heterogeneity in multiple myeloma revealed by multi-region sequencing. Nat Commun 8, 268 (2017).

30. Papadea, C., Reimer, C. B. & Check, I. J. IgG subclass distribution in patients with multiple myeloma or with monoclonal gammopathy of undetermined significance. Ann Clin Lab Sci 19, 27–37 (1989).

31. Zhang, L., Fok, J. H. L. & Davies, F. E. Heat shock proteins in multiple myeloma. Oncotarget 5, 1132–1148 (2014).

32. Caillot, M., Dakik, H., Mazurier, F. & Sola, B. Targeting Reactive Oxygen Species Metabolism to Induce Myeloma Cell Death. Cancers 13, 2411 (2021).

33. Tellier, J. & Nutt, S. L. Standing out from the crowd: How to identify plasma cells. European Journal of Immunology 47, 1276–1279 (2017).

34. Liu, M. et al. S100 Calcium Binding Protein Family Members Associate With Poor Patient Outcome and Response to Proteasome Inhibition in Multiple Myeloma. Front. Cell Dev. Biol. 9, (2021).

35. Wang, G., Fan, F., Sun, C. & Hu, Y. Looking into Endoplasmic Reticulum Stress: The Key to Drug-Resistance of Multiple Myeloma? Cancers (Basel) 14, 5340 (2022).

36. Almanza, A. et al. Endoplasmic reticulum stress signalling - from basic mechanisms to clinical applications. FEBS J 286, 241–278 (2019).

37. Dolina, J. S., Van Braeckel-Budimir, N., Thomas, G. D. & Salek-Ardakani, S. CD8+ T Cell Exhaustion in Cancer. Front Immunol 12, 715234 (2021).

38. Joshua, D. E. et al. Treg and Oligoclonal Expansion of Terminal Effector CD8+ T Cell as Key Players in Multiple Myeloma. Front. Immunol. 12, (2021).

39. Bai, Y., Hu, M., Chen, Z., Wei, J. & Du, H. Single-Cell Transcriptome Analysis Reveals RGS1 as a New Marker and Promoting Factor for T-Cell Exhaustion in Multiple Cancers. Front. Immunol. 12, (2021).

40. Bian, F. et al. Spatial Transcriptomics Reveals the Requirement of ADGRG6 in Maintaining Chondrocyte Homeostasis in Mouse Growth Plates. bioRxiv (2023). 10.1101/2023.09.21.558739.

41. Zheng, X. et al. A single-cell and spatially resolved atlas of human osteosarcomas. J Hematol Oncol 17, 71 (2024).

42. Maas, R. R. et al. The local microenvironment drives activation of neutrophils in human brain tumors. Cell 186, 4546–4566.e27 (2023).

43. Wang, W. et al. Identification and Validation of a Novel RNA-Binding Protein-Related Gene-Based Prognostic Model for Multiple Myeloma. Front. Genet. 12, (2021).

44. Boiarsky, R. et al. Single cell characterization of myeloma and its precursor conditions reveals transcriptional signatures of early tumorigenesis. Nat Commun 13, 7040 (2022).

45. Samur, M. K., Szalat, R. & Munshi, N. C. Single-cell profiling in multiple myeloma: insights, problems, and promises. Blood 142, 313–324 (2023).

46. Lamanuzzi, A. et al. Thrombopoietin Promotes Angiogenesis and Disease Progression in Patients with Multiple Myeloma. Am J Pathol 191, 748–758 (2021).

47. van Buul, J. D. & Hordijk, P. L. Signaling in Leukocyte Transendothelial Migration. Arteriosclerosis, Thrombosis, and Vascular Biology 24, 824–833 (2004).

48. Fu, J. et al. The checkpoint inhibitor PD-1H/VISTA controls osteoclast-mediated multiple myeloma bone disease. Nat Commun 14, 4271 (2023).

49. Rustad, E. H. et al. Baseline identification of clonal V(D)J sequences for DNA-based minimal residual disease detection in multiple myeloma. PLOS ONE 14, e0211600 (2019).

50. Space Ranger - Official 10x Genomics Support. 10x Genomics https://www.10xgenomics.com/support/software/space-ranger/latest.

51. Stuart, T. et al. Comprehensive Integration of Single-Cell Data. Cell 177, 1888–1902.e21 (2019).

52. Bergenstråhle, J., Larsson, L. & Lundeberg, J. Seamless integration of image and molecular analysis for spatial transcriptomics workflows. BMC Genomics 21, 482 (2020).

53. Hafemeister, C. & Satija, R. Normalization and variance stabilization of single-cell RNA-seq data using regularized negative binomial regression. Genome Biology 20, 296 (2019).

54. Ma, Y. & Zhou, X. Spatially informed cell-type deconvolution for spatial transcriptomics. Nat Biotechnol 40, 1349–1359 (2022).

55. Kassambara, A. & Mundt, F. factoextra: Extract and Visualize the Results of Multivariate Data Analyses. (2020).

56. Taiyun. taiyun/corrplot. (2023).

